# RbAp46/48^LIN-53^ and HAT-1 are required for initial CENP-A^HCP-3^ deposition and de novo centromere formation in *Caenorhabditis elegans* embryos

**DOI:** 10.1101/2020.04.12.019257

**Authors:** Zhongyang Lin, Karen Wing Yee Yuen

## Abstract

Foreign DNA microinjected into the *Caenorhabditis elegans* germline forms episomal extra-chromosomal arrays, or artificial chromosomes (ACs), in embryos. Injected linear, short DNA fragments concatemerize into high molecular weight (HMW)-DNA arrays that are visible as punctate DAPI-stained foci in oocytes, which undergo chromatinization and centromerization in embryos. The inner centromere, inner and outer kinetochore components, including AIR-2, CENP-A^HCP-3^, Mis18BP1^KNL-2^ and BUB-1, assemble onto the nascent ACs during the first mitosis. Yet, due to incomplete DNA replication of the nascent ACs, centromeric proteins are not oriented at the poleward faces of the nascent ACs in mitosis, resulting in lagging ACs. The DNA replication efficiency of ACs improves over several cell cycles. We found that a condensin subunit, SMC-4, but not the replicative helicase component, MCM-2, facilitates *de novo* CENP-A^HCP-3^ deposition on nascent ACs. Furthermore, H3K9ac, H4K5ac, and H4K12ac are highly enriched on newly chromatinized ACs. HAT-1 and RbAp46/48^LIN-53^, which are essential for *de novo* centromere formation and segregation competency of nascent ACs, also hyperacetylate histone H3 and H4. Different from centromere maintenance on endogenous chromosomes, where Mis18BP1^KNL-2^ functions upstream of RbAp46/48^LIN-53^, RbAp46/48^LIN-53^ depletion causes the loss of both CENP-A^HCP-3^ and Mis18BP1^KNL-2^ initial deposition at *de novo* centromeres on ACs.

## INTRODUCTION

Histone H3 variant, CENP-A, replaces histone H3 in the centromeric nucleosomes and serves as the foundation for building the kinetochore, which connects the sister chromatids to opposite spindles and orchestrates chromosome movement. Centromere propagation through cell cycles and generations is crucial for ensuring accurate chromosome segregation and maintenance of genome integrity, which relies on histone chaperones to deposit CENP-A precisely to the centromeric regions of sister chromatids.

Ectopic formation of a centromere can cause dicentric chromosome formation, which undergoes chromosome breakage-fusion cycle, leading to chromosomal rearrangements, chromosome loss or gain, aneuploidy, and potentially chromosome instability and tumorigenesis. Many cases of neocentromeres were found in human patients with congenital abnormalities or developmental disorders (1). The mechanism of new centromere formation is still not fully understood because of the technical challenges in tracing the early events of neocentromere formation in patients’ cells. However, this phenomenon has been observed in diverse species. For example, tethering CENP-A-specific chaperones to a euchromatin locus or overexpressing CENP-A could cause ectopic CENP-A localization and ectopic centromere formation (2,3). Besides, transforming or transfecting centromeric DNA into yeast or human cells can form artificial chromosomes with *de novo* centromeres (2,3). However, artificial chromosome formation in these species often relies on the presence of their own centromeric DNA sequences, requiring long-term drug selection, and has relatively low frequencies of *de novo* centromere formation, which has limited their applications in studying the early events of new centromere formation.

In *C. elegans*, injecting foreign DNA, even devoid of *C. elegans* sequences, into its gonad could form episomal extra-chromosomal arrays, also known as artificial chromosomes (ACs), in the embryonic cells. These ACs can be propagated mitotically and inherited through subsequent generations (4,5). These heritable ACs have established a functional holocentromere, rather than hitchhiking on the endogenous chromosomes (6). Dissecting the mechanism of *de novo* centromere establishment on ACs could help to understand the process of neocentromere formation on endogenous chromosomes.

In the present study, after injection of short, linear DNA, we investigated the timing of *de novo* CENP-A^HCP-3^ deposition on ACs, and demonstrated that CENP-A^HCP-3^ starts to assemble on ACs after fertilization. Another inner kinetochore protein, Mis18BP1^KNL-2^ and an inner centromere protein, AIR-2, are also recruited to the nascent ACs in the first mitosis. The ACs attempt to segregate in the first cell division, but with anaphase bridges. We also analyzed the histone post-translational modifications (PTMs) that co-occur with *de novo* CENP-A^HCP-3^ deposition on nascent ACs in one-cell embryos. Based on the profiles of the enriched histone PTMs on nascent ACs, we depleted the relevant histone modifiers or the associated histone chaperones by RNA interference (RNAi), and analyzed the AC segregation rate by live-cell imaging and the centromeric protein signals by immunofluorescence analysis. We demonstrated that HAT-1 and RbAp46/48^LIN-53^ are required for the enriched H3K9ac, H4K5ac and H4K12ac on nascent ACs in one-cell embryos. Depleting HAT-1, RbAp46/48^LIN-53^ or both will reduce or abolish ACs’ segregation competency by reducing *de novo* CENP-A^HCP-3^ deposition on nascent ACs. Surprisingly, at the *de novo* centromere on ACs, RbAp46/48^LIN-53^ depletion leads to the loss of both CENP-A^HCP-3^ and Mis18BP1^KNL-2^ initial deposition, which suggests that while Mis18BP1^KNL-2^ could be the self-directing factor for centromere maintenance in existing centromeres, it is downstream of RbAp46/48^LIN-53^ in initial centromere establishment. We show that efficient *de novo* CENP-A^HCP-3^ deposition on ACs also requires condensin subunit SMC-4, but it is independent of DNA replicative helicase component, MCM-2. These results demonstrate that the mechanism of *de novo* CENP-A^HCP-3^ deposition on ACs requires histone acetyltransferase HAT-1, CENP-A deposition machinery, including histone chaperone RbAp46/48^LIN-53^, together with M18BP1^KNL-2^ and SMC-4.

## MATERIAL AND METHODS

### Worm strains and maintenance

The CRISPR/Cas9 transgenic technique described by Dickinson and Goldstein (7) was used to design and generate a GFP-tagged HAT-1 at the endogenous locus. PCR genotyping was done using primer set: Seq-Hat-1 (Table S1). Worm strains used in this study are listed in Table S2. All worms were maintained at 22°C on standard EZ plates seeded with *E. coli* OP50.

### Double-stranded RNA (dsRNA) synthesis and RNA interference (RNAi)

PCR primers were designed to amplify a region of target genes from N2 *C. elegans* genomic DNA or cDNA. T3 promoter (AATTAACCCTCACTAAAGG) or T7 promoter (TAATACGACTCACTATAGG) was added to the 5’ end of primers. Primers (Table S1) were selected using NCBI-Primer-Blast and were subjected to BLAST search using the *C. elegans* genome to confirm the primer specificity. PCR was performed using TaKaRa Ex Taq® DNA Polymerase and the PCR products were purified by Qiagen PCR purification kit. Purified PCR products were subjected to *in vitro* transcription using Ambion T3 and T7 MEGAscript® Kit at 37°C for 4-6 hours. Reaction products were digested with TURBO DNase at 37 °C for 15 min and purified using Ambion MEGAclear™ Kit. Eluates were incubated at 68°C for 10 minutes followed by 37°C for 30 minutes for complementary RNA annealing. Annealed dsRNA was adjusted to 1 μg/μL in ddH_2_O for microinjection. For RNAi, L4 hermaphrodites were injected with dsRNA (1 μg/μL) and recovered at 22°C for 24 hours before further analysis. For RNAi plus AC introduction, L4 hermaphrodites were injected with dsRNA (1 μg/μL) and recovered at 22°C for 18 hours to reach the young adult stage. The RNAi-treated worms were then injected with p64xLacO plasmid DNA, linearized by AfaI (L64×LacO) and purified, into the gonad and recovered at 22°C for another 5 hours before live imaging or Immunofluorescence staining.

### Live cell imaging and AC segregation assay

Episomal artificial chromosomes (ACs) were visualized by injecting DNA containing LacO tandem repeats as reported previously (6), except that we used linear DNA (L64×LacO) for microinjection. Injected worms were recovered on OP50-seeded plates for 5-8 hours after microinjection. 3-4 worms were then dissected in 2 μl M9 buffer to release embryos. Embryos were mounted on a freshly prepared 2% agarose pad and the slide edges were sealed with Vaseline. Live-cell images were taken with a Carl Zeiss LSM710 laser scanning confocal microscope with a 16 AC Plan-Neofluar 40x Oil objective lens and PMT detectors. Stacks with 17×1.32 μm planes were scanned for each embryo in a 3x zoom and a 1-minute or 30 second time interval, with 3.15 μs pixel dwell and 92 μm pinhole. Laser power for 488 nm and 543 nm was set at 5.5% and 6.5%, respectively.

To determine the AC segregation rates, every dividing cell that contains at least one AC was counted as one sample. Each division was categorized as either containing at least a segregating AC or containing all non-segregating AC(s). Segregating ACs were defined as those that aligned with the metaphase plate and segregated with endogenous chromosomes during anaphase. Non-segregating ACs were defined as those that remained in the cytoplasm or nucleus and did not segregate in mitosis. The AC segregation rate was calculated as the number of dividing cells containing segregating ACs over the total number of dividing cells containing ACs. Among the segregating ACs, those with anaphase bridges were referred to as ACs that attempted to segregate, but ACs were lagging during anaphase, and the AC segregation process is incomplete.

### Immunofluorescence (IF) staining

Embryos were freeze-cracked after dissection of adult worms and fixed in −20°C methanol for 30 minutes. Embryos were then rehydrated in PBS [137 mM NaCl, 2.7 mM KCl, 4.3 mM Na_2_HPO_4_, 1.4 mM KH_2_PO_4_] for 5 minutes and blocked by AbDil [4% BSA, 0.1% Triton-X 100 in PBS] at room temperature for 20 minutes. Primary antibody incubation, using rabbit (Rb)-anti-HCP-3 (1:1000; Novus biologicals 29540002), Rb-anti-KNL-2 (1:500, a gift from Desai Lab), Rb-anti-SMC-4 (1:500, a gift from Desai Lab), Mouse (Ms)-anti-LacI (1:250, Millipore 05-503), Rb-anti-H3K9ac (1:500; Millipore ABE18), Rb-anti-H4K5ac (1:500, Abcam ab51997), Rb-anti-H4K8ac (1:500, Abcam ab45166), Rb-anti-H4K12ac (1:500, Abcam ab177793), Rb-anti-H4K16ac (1:500, Abcam ab109463), Rb-anti-H3K27me3 (1:500; Millipore 07-449), Rb-anti-H3K4me (1:500, Abcam ab176877), Rb-anti-H3K4me2 (Novus Biologicals NB21-1022), Rb-anti-H3K4me3(1:500, Abcam ab8580), Ms-anti-H3K9me2 (1:500 Abcam, ab1220), Rb-anti-H3K9me3 (1:500, Abcam Ab8898), Rb-anti-H3 (1:1000, Abcam Ab18521) or Rb-anti-H4 (1:1000, Ab10158) was performed at 4°C overnight. Slides were washed with PBST 3×10 minutes. The slides were then incubated with goat-anti-Ms-IgG FITC (1:100,000; Jackson ImmunoResearch Laboratories, 115-096-062) and goat-anti-Rb-IgG Alexa 647-conjugated secondary antibody (1:100,000; Jackson ImmunoResearch Laboratories, 111-606-045) at room temperature for 1 hour, followed by DAPI (1μg/mL) staining for 15 minutes. The fluorescent signal of mChery::H2B is detectable after methanol fixation and was measured without antibody incubation. Mounting was done using ProLong gold antifade reagent (Life Technologies). Images were acquired from Zeiss LSM 780 upright confocal microscope with a Plan-Apochromat 40×1.4 Oil DIC M27 objective and PMT detectors. Embryos were captured as z stacks with a z-step size at 0.4 μm and 3.15 μs of pixel dwell time. Stacks with 30-35 × 0.4 μm planes were scanned for each embryo in a 4x zoom. DAPI, FITC and Alex647 channels were scanned with 32 μm pinhole, and the images were saved in 16 bits format.

### 5-Ethylnyl-2’-deoxyuridine (EdU) staining of one-cell stage embryos

L4 worms were grown on *perm-1* dsRNA-expressing bacteria diluted 1/6 with OP50 for 24 hours (8). EdU staining of embryos for 15 minutes was performed as described previously (9).

### Image quantification

Images were processed with Fiji 2.0.0. For immunofluorescence, 31 z-sections were acquired with a spacing of 0.4 µm for each embryo. The region of interest (ROI) and the number of z-stacks for each target object were manually selected. A larger area enclosing the whole ROI within the embryo was drawn in each sample (ROI-L). Integrated density (IntDen) equals to area times mean grey value. For each channel, the integrated density of the ROI and ROI-L from all selected z-stacks containing the target object were summed (ROI^IntDen^ and ROI-L^IntDen^). The area between ROI and ROI-L were used for calculating the mean grey value of background following the equation Bg^mean^ = (ROI-L^IntDen^ - ROI^IntDen^)/ (ROI-L^area^ - ROI^area^). The corrected integrated density of each targeted protein, histone modification or EdU in ROI = ROI^IntDen^ – (ROI^area^ x Bg^mean^) was then normalized with the corrected integrated density of DAPI.

## RESULTS

### Formation of artificial chromosomes (ACs) through chromatinization and *de novo* centromerization of foreign DNA in *C. elegans* one-cell embryos

To follow the fate of foreign DNA injected into the syncytial germline of *C. elegans*, the germline, oocytes and embryos were imaged 5 hours after injection of linearized 64 copy-LacO arrays (L64×LacO). To identify the timing of DNA array formation, DAPI (4’, 6-diamidino-2-phenylindole) staining was used to indicate the location, size and morphology of the concatemerized injected foreign DNA in the germline. To determine the timing of nucleosome assembly and *de novo* centromere formation, live-cell imaging of histone H2B (H2B::mCherry) and CENP-A^HCP-3^::GFP was used to indicate the status of canonical histone deposition and centromeric nucleosome assembly, respectively. DAPI staining of the HMW foreign DNA could not be observed in the syncytial gonad where L64×LacO was injected. However, punctate DAPI foci were found in the cytoplasm of the diplotene and diakinesis oocytes, suggesting that the injected DNA fused into high molecular weight (HMW), extra-chromosomal DNA arrays (Figure 1A and Figure S1 A). Based on the morphology and size, the HMW DNA arrays can be easily distinguished from the 6 highly compacted endogenous bivalent chromosomes. These DNA arrays in oocytes lack histone H2B, CENP-A^HCP-3^ and Mis18BP1^KNL-2^ in oocytes (Figure 1A and Figure S1A). However, in fertilized zygotes, these DNA arrays became artificial chromosomes (ACs) that contain detectable histone H2B (6) and CENP-A^HCP-3^ (Figure 1B), indicating that chromatinization and *de novo* centromerization of foreign DNA have begun in one-cell embryos after fertilization. CENP-A^HCP-3^ signal was observed on ACs as early as in embryos undergoing meiosis I (Figure 1B). Live-cell imaging showed that newly formed ACs aligned at the metaphase plate and attempted to segregate during the first mitosis (Figure 1C, S1B and Suppl.Video1). All of the aligned ACs were pulled towards opposite poles at anaphase, but all form chromosome bridges (Figure 1C and S2G). This result suggests that the kinetochore on ACs were sufficient to attach to the mitotic spindles. To elucidate the cause of chromosome bridge formation, we investigate the level and orientation of kinetochore proteins on these newly formed ACs.

**Figure 1.**
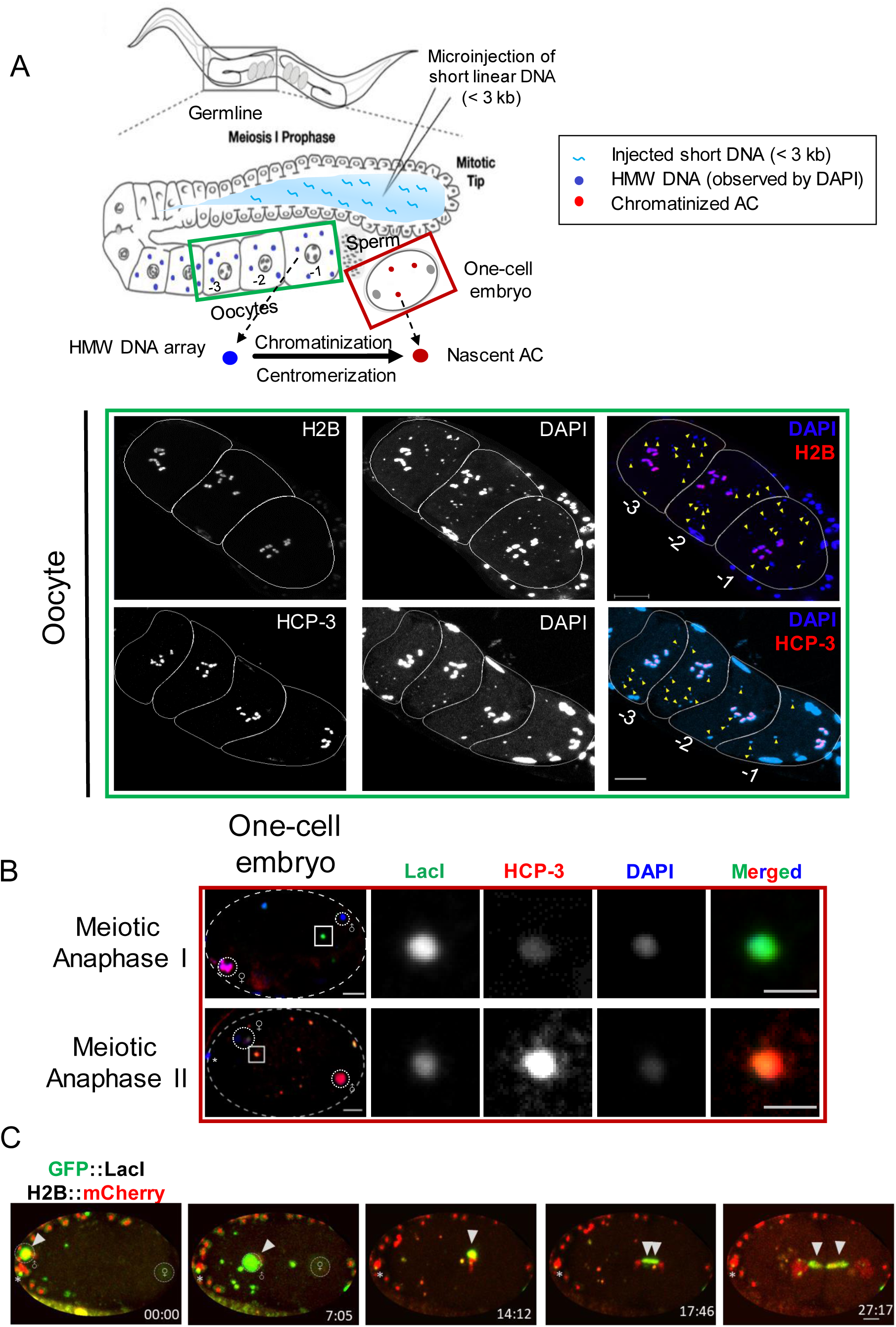
Chromatinization and *de novo* CENP-A^HCP-3^ formation on foreign HMW DNA arrays to form artificial chromosomes (ACs) in fertilized one-cell embryos. (A) A schematic diagram of delivering short, linearized p64xLacO plasmid (L64×LacO) DNA into *C. elegans* gonad by microinjection. DAPI stained six condensed bivalent endogenous chromosomes and the HMW DNA arrays concatemerized from the injected foreign DNA. Representative immunofluorescence images of the H2B::mCherry and of CENP-A^HCP-3^ on bivalent chromosomes in oocytes with multiple DAPI foci. Yellow arrowheads indicate the HMW foreign DNA arrays. Scale bar represents 10 μm. (B) Representative immunofluorescence images show that nascent artificial chromosomes (ACs) assembled from the HMW DNA arrays contain detectable CENP-A^HCP-3^ signals in one-cell embryos at meiosis I and II, respectively. White dash ovals show the paternal and maternal DNA, and * represents the polar body. Scale bar represents 5 μm. A higher-magnification view of the representative ACs (white square) is shown on the right, in which the scale bar represents 2 μm for the magnified images. (C) Time-lapse images following an AC, which was attempting to segregate during the first mitosis in one-cell embryos. The time-lapse after fertilization was shown (mm:ss). Scale bar represents 5 μm.

### Impaired DNA replication does not affect *de novo* CENP-A deposition, but causes centromere disorganization on metaphase ACs

MCM-2, a component of the MCM2-7 replicative helicase, is essential for DNA replication (10), and it is also a histone chaperone for restoring histones to newly synthesized DNA (11). Because the formation of the MCM-2-7 complex depends on each of its subunits, depleting MCM-2 will prevent MCM-2-7 complex assembly and block the process of DNA replication (10). To determine the effect of MCM-2 on initial CENP-A^HCP-3^ deposition and *de novo* centromere formation, we performed double-stranded RNA (dsRNA) microinjection to deplete *mcm-2* mRNA. None of the embryos from the injected worms were able to hatch (data not shown), suggesting that the RNAi of *mcm-2* is highly efficient. However, our result shows that MCM-2 depletion does not prevent ACs from aligning at the metaphase plate and attempting to segregate with the endogenous chromosomes (Figure S2H and S2I). Immunofluorescence analysis shows that the essential inner kinetochore proteins, CENP-A^HCP-3^ and Mis18BP1^KNL-2^, were both present on the nascent ACs that lined up at metaphase plate and on the bridging ACs at anaphase in one-cell embryos (Figure 2A and 2B). Quantification of the integrated intensity of CENP-A^HCP-3^ on nascent ACs, normalized to the total amount of DNA on ACs, shows no significant reduction of CENP-A^HCP-3^ incorporation in *mcm-2* RNAi-treated embryos, which suggests that *de novo* CENP-A^HCP-3^ deposition is independent of MCM-2 (Figure 2E and 2F). Moreover, the recruitment of outer kinetochore and spindle checkpoint component, BUB-1, on ACs is also not abolished upon MCM-2 depletion (Figure S2J).

**Figure 2.**
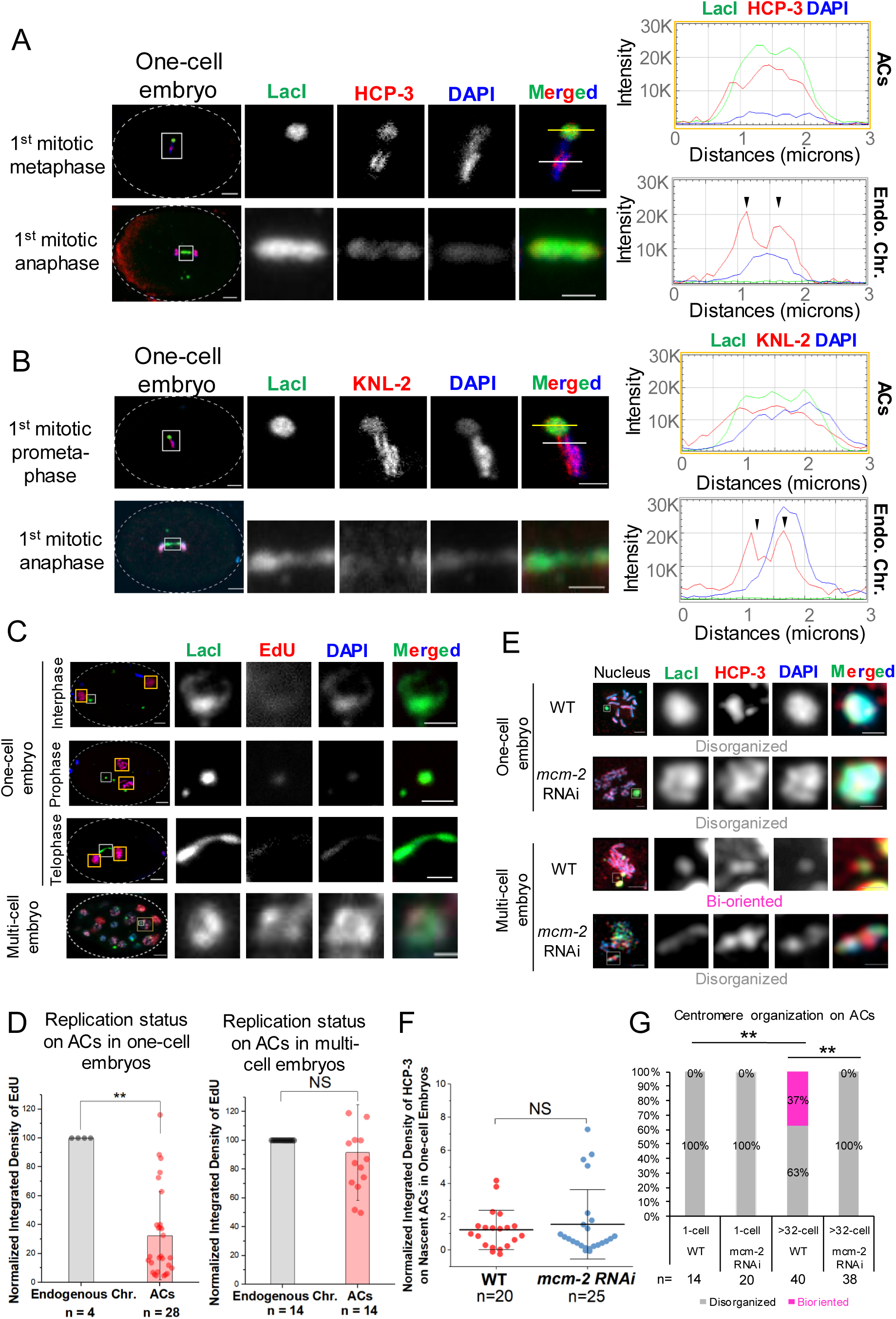
Impaired DNA replication causes centromere disorganization on ACs and anaphase bridges. (A & B) Immunofluorescence staining of ACs (LacI), inner kinetochore proteins, CENP-A^HCP-3^ (A) or M18BP1^KNL-2^ (B), and chromatin (DAPI) at metaphase and anaphase in one-cell embryos. The line-scan analysis shows the signal intensity of CENP-A^HCP-3^ and M18BP1^KNL-2^, respectively, on the metaphase plate of ACs and endogenous chromosomes. Scale bar represents 5 μm. A 3-μm line was drawn across the metaphase plate in the high magnification panels, and the signal intensities were measured (Yellow line: AC; White line: Endogenous chromosomes). Scale bar in magnified panels represents 2 μm. The plot shows signal intensities from each channel along the line. Green line: LacI; Red line: CENP-A^HCP-3^ (A) or M18BP1^KNL-2^ (B); Blue line: DAPI. The black arrowheads indicate the poleward orientation of CENP-A^HCP-3^ on endogenous chromosomes. CENP-A^HCP-3^ on the AC lacks such bi-orientation at metaphase. (C) EdU staining of nascent ACs in one-cell embryos at interphase, prophase and telophase, respectively, and ACs in a multi-cell embryo. (D) Comparison of the average uptake of EdU after 15 minutes of incubation (normalized to DAPI) on endogenous chromosomes and nascent ACs in mitotic one-cell and multi-cell embryos, respectively. All ACs or endogenous chromosomes in one-cell or multi-cell embryos were pooled together (n = number of samples) for calculating the mean of EdU integrated density. The bar chart shows the mean EdU signal on ACs relative to that on endogenous chromosomes. The error bars represent standard deviation (SD). Significant differences are analyzed by the Student t-test (**, p < 0.01; NS, not significant). (E) Immunofluorescence of CENP-A^HCP-3^ on ACs in untreated wild-type (WT) or *mcm-2* RNAi-treated one- and multi-cell stage embryos during prometaphase. Scale bar represents 2 μm. CENP-A^HCP-3^ on the entire AC is described as “disorganized”, while CENP-A^HCP-3^ on the poleward sides of the AC is described as “bi-oriented”. (F) A scatter plot shows the quantification of integrated density of CENP-A^HCP-3^ signal on ACs in WT and in *mcm-2* RNAi-treated one-cell embryos. The error bars represent standard deviation (SD). Significant differences are analyzed by the Student t-test (NS, not significant). (G) Quantification of the percentage of ACs with disorganized or bi-oriented CENP-A^HCP-3^ on AC in one- and multi-cell stage WT or *mcm-2* RNAi-treated embryos. The number of ACs (n) analyzed was indicated. Significant differences are analyzed by the Fisher’s exact test (**, p < 0.01).

Proper sister chromatid segregation depends on the bi-orientation of sister kinetochores on sister chromatids, and the capture of microtubules emanating from opposite centrosomes. However, the initially formed kinetochore on nascent ACs lacks bi-orientation at metaphase (Figure 2A), but is all over the AC, potentially causing merotelic attachments of microtubules to the nascent ACs, which leads to the lagging of ACs during chromosome segregation in anaphase. Because inner centromeric proteins, such as AIR-2 and condensin II, also contribute to forming bi-oriented kinetochores (12), we checked if inner centromere proteins are present on nascent ACs that aligned at the metaphase plate. Our immunofluorescence analysis shows that inner centromeric protein, AIR-2, and condensin II component, SMC-4, are both recruited to the nascent ACs at the metaphase plate (Figure S2A and S2B). In the properly segregated endogenous chromatids, SMC-4 dissociates from the chromatin during anaphase. In contrast, SMC-4 is still found in the center of the AC chromatin bridges in late anaphase (Figure S2B). DNA replication has been shown to be needed for chromatin decondensation in *C. elegans* embryos in anaphase (9), which is consistent with the loss of condensin II component SMC-4 during anaphase. We found that impairing DNA replication by hydroxyurea (HU) treatment also results in the persistent presence of SMC-4 on the bridging endogenous chromosomes in late anaphase (Figure S2C). We proposed that incomplete DNA replication is the reason why nascent ACs are lagging in anaphase.

We then measured the DNA replication efficiency on ACs by their 5-ethynyl-2’-deoxyuridine (EdU) incorporation efficiency in one-cell stage and multi-cell stage embryos. In one-cell embryos, ACs showed at least 68% reduction of EdU incorporation when compared to endogenous chromosomes (Figure 2C and D), which suggests that DNA replication is less efficient on ACs in early-stage embryos. Also, live-cell imaging shows that MCM-4::mCherry (another component of MCM-2-7 DNA replicative helicase complex), which is supposed to dissociate from endogenous chromosomes before metaphase, still has prolonged association with the ACs (Figure S2D), indicating that DNA replication on ACs has yet to be completed even at metaphase. Thus, we proposed that incomplete DNA replication on ACs (Figure 2C) may contribute to the lack of AC’s kinetochore bi-orientation and the lagging ACs in early-stage embryos. In multi-cell embryos, however, we found that the replication of ACs has improved, where the EdU incorporation rate on ACs is comparable to that on endogenous chromosomes (Figure 2C and 2D). Consistently, we found more ACs in multi-cell embryos (37%) possess bi-orientated centromeres than in one-cell embryos (0%) in wild-type (Figure 2E and 2G). As expected, more ACs segregated evenly in multi-cell stage embryos than in one-cell stage embryos (Figure S2F and S2G).

### Condensin II facilitates *de novo* CENP-A^**HCP-3**^ **deposition on nascent ACs in one-cell embryos**

In *C. elegans*, condensin II complex co-localizes with centromere proteins on metaphase chromosomes (13,14), and is proposed to have a specific function at the centromere in addition to chromatin condensation. In human cells and *Xenopus* egg extracts, condensin II is required for new CENP-A deposition in mitotic cells and new CENP-A loading in 1^st^ mitosis, respectively (15,16). As the SMC-4 signal is positive on nascent ACs, we further determined if condensin II contributes to *de novo* centromere formation on ACs in *C. elegans* embryos. Quantification of the CENP-A^HCP-3^ level on ACs shows a significant reduction of CENP-A^HCP-3^ level in *smc-4* RNAi-treated embryos, suggesting that condensin II facilitates *de novo* CENP-A^HCP-3^ deposition on nascent ACs in *C. elegans* embryos (Figure 3).

**Figure 3.**
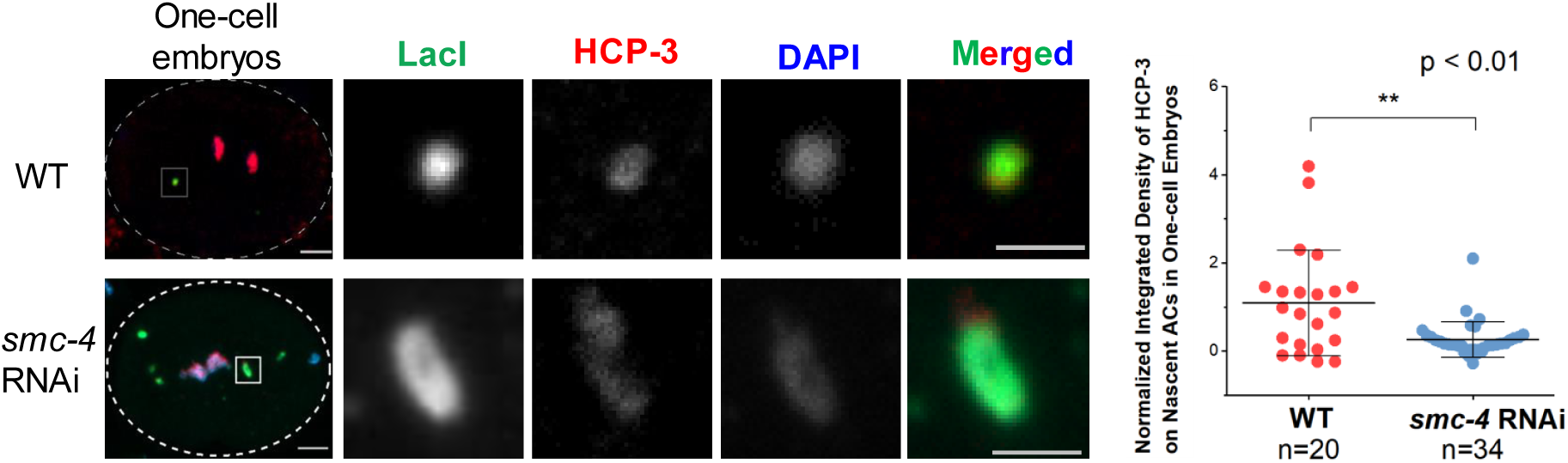
Depletion of condensin II subunit, SMC-4, reduces *de novo* CENP-A^HCP-3^ deposition on ACs. Immunofluorescence of CENP-A^HCP-3^ on ACs in WT and *smc-4* RNAi-treated one-cell embryos. Embryos were stained with antibodies against LacI (green), CENP-A^HCP-3^ (red) and DAPI (blue). Scale bar represents 5 μm. A higher-magnification view of the ACs (white square) is shown on the right. Scale bar represents 2 μm for the magnified images. A scatter plot shows the quantification of normalized integrated density of CENP-A^HCP-3^ signal on ACs in WT and *smc-4* RNAi-treated one-cell embryos. The integrated density was normalized with that of DAPI. The number of samples (n) analyzed was indicated. The number of samples (n) analyzed was indicated. The error bars represent SD. Significant differences are analyzed by the Student’s t-test, **, p < 0.01.

### The spectrum of histone post-translational modifications (PTMs) on nascent ACs

To further identify essential factors in *de novo* centromere formation, we profiled the histone PTMs on the nascent ACs. We hypothesize that histone-modifying codes that co-exist with chromatinization and centromerization on newly formed ACs in one-cell embryos may help us identify the required factors. We chose several histone PTMs that have been reported to be associated with centromere function. The spectrum of histone PTMs on newly formed ACs in one-cell embryos by immunofluorescence analysis (Figure 4 and S3) is summarized in Table 1. For PTMs that are associated with transcription activity, we analyzed H3K4me1, H3K4me2 and H3K4me3 on newly formed ACs (Figure 4A). Methylation of H3K4 is associated with active transcription. The presence of a medium level of H3K4me1 on ACs, even with the lack of H3K4me2 and H3K4me3, suggests that ACs might be actively transcribing. It has been shown that RNAPII docking facilitates *de novo* centromere formation in ACs formed by injecting circularized, supercoiled p64xLacO plasmid (17). Consistently, the signal intensities of H4K5ac (Figure 4B), H4K12ac (Figure 4C), H3K9ac (Figure 4D) and H4K20me (Figure 4E) on nascent ACs are significantly higher than that on endogenous chromosomes, with 3-, 3.5-, 2.2- and 10-fold enrichment, respectively. Meanwhile, the DNA replication-associated histone PTM, H3K56ac, on nascent ACs has dimmer signal intensity as compared to that on endogenous chromosomes (Figure 4A and S3). This is consistent with the above finding that replication is less efficient on nascent ACs than on endogenous chromosomes (Figure 2C and 2D). Moreover, heterochromatin-associated histone PTMs, including H3K9me2, H3K9me3 and H3K27me3, are undetectable on nascent ACs in one-cell embryos (Figure 4A and S3), consistent with our previous finding that heterochromatin is dispensable for *de novo* centromere formation (6).

**Table 1.** Summary of the profile of histone PTMs on newly formed ACs in one-cell embryos.

| CENP-A and histone PTMs | Signal intensity on newly formed ACs* | Function in Centromere | Associated histone modifiers or factors | Reference on function and associated factors |
| --- | --- | --- | --- | --- |
| CENP-A<sup>HCP-3</sup> | ++ | Epigenetic mark of centromere | M18BP1<sup>KNL-2</sup> & RbAp46/48<sup>LIN-53</sup> | (61) |
| H4K20me | +++ | CENP-A nucleosome | SET-1 (18) | (38,62) |
| H4K5ac | +++ | Newly synthesized histone H4 | RbAp46/48<sup>LIN-53</sup> & HAT-1 | (48,63) |
| H4K12ac | +++ | Newly synthesized histone H4 | RbAp46/48<sup>LIN-53</sup> & HAT-1 | (48,63) |
| H3K9ac | +++ | Newly synthesized histones/ open chromatin marker | RbAp46/48<sup>LIN-53</sup>, HAT-1 & unknown factors | (40) |
| H3K56ac | + | Newly synthesized histones deposited during DNA replication | MCM-2 & ASF-1 | (11,64) |
| H3K4me | + | Permissive transcription | SET-17 | (65,66) |
| H3K4me2 | - | Active transcription | SET-17 & SET-30 | (66) |
| H3K4me3 | - | Robust transcription | ASH-2 | (67) |
| H3K9me2 | - | Heterochromatin mark | MET-2 | (68) |
| H3K9me3 | - | Heterochromatin mark | MES-2 | (68) |
| H3K27me3 | - | Heterochromatin mark | MES-2 | (65) |

+++ : signal on ACs is significantly higher than that on endogenous chromosomes
++ : signal on ACs is comparable to that on endogenous chromosomes
+ : signal on ACs is significantly lower than that on endogenous chromosomes
- : signal on ACs is undetectable

**Figure 4.**
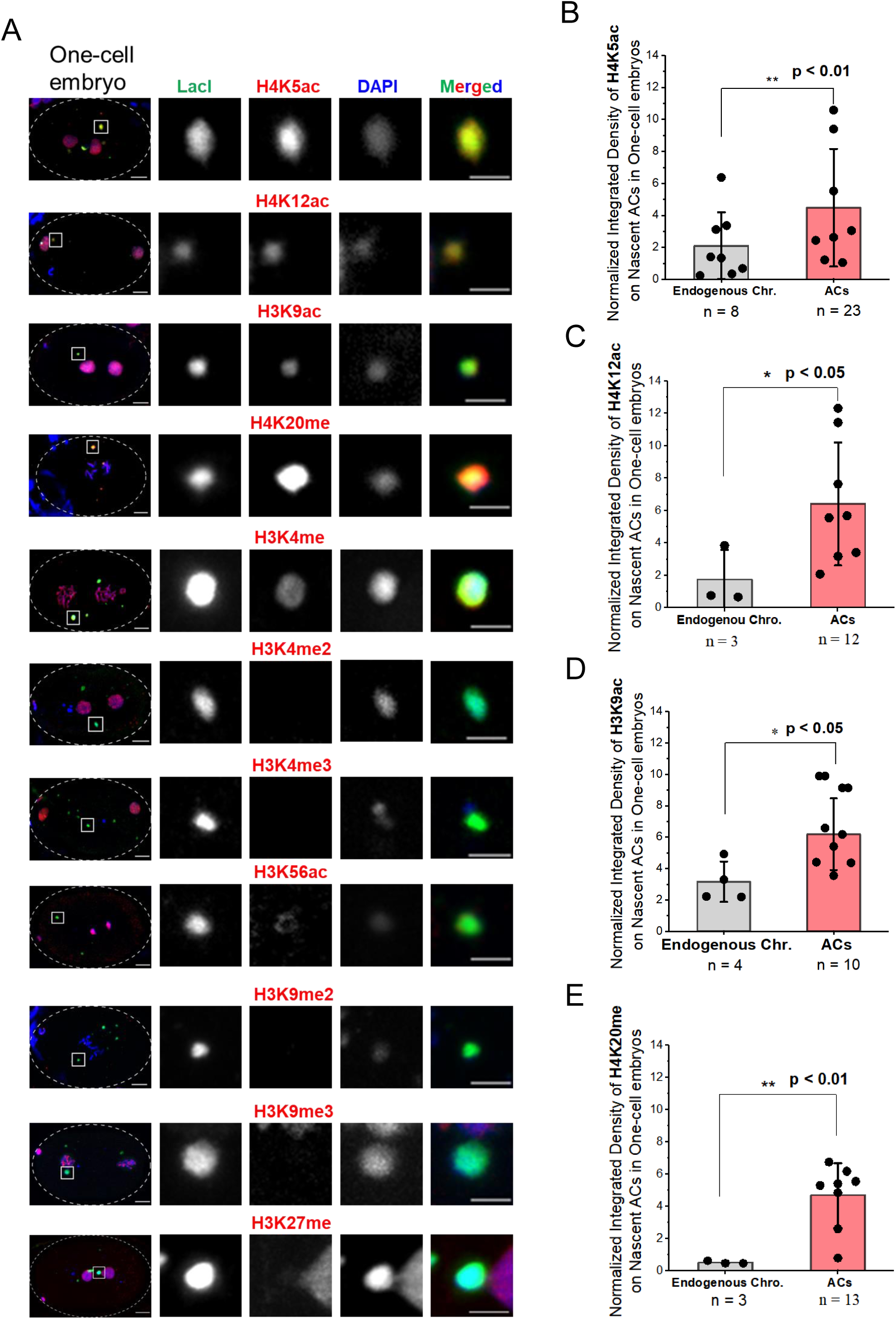
Profiling of histone post-translational modifications (PTMs) on nascent ACs in one-cell embryos by immunofluorescence (IF) staining. (A) Representative immunofluorescence images of H4K5ac, H4K12ac, H3K9ac, H4K20me, H3K4me, H3K4me2, H3K4me3, H3K56ac, H3K9me2, H3K9me3 and H3K27me3 on endogenous chromosomes and newly formed ACs in one-cell embryos. Embryos were stained with antibody against LacI (green), antibodies against a histone PTM (red) and DAPI (blue). Scale bar represents 5 μm. A higher-magnification view of the ACs (white square) is shown on the right. Scale bar represents 2 μm for the magnified images. The representative images were contrast adjusted. The box plot shows the quantification result of the normalized integrated density of (B) H4K5ac; (C) H4K12ac; (D) H3K9ac; or (E) H4K20me signal on endogenous chromosomes and on ACs in one-cell embryos. Only quantifications of the enriched PTMs are shown. Other PTM levels are shown in Table 1. For quantification of PTMs on ACs and endogenous chromosomes, the signal density of each PTM was normalized with that of DAPI. The number of samples (n) analyzed was indicated. The error bars represent SD. Significant differences are analyzed by the Student t-test (*, p < 0.05; **, p < 0.01).

### The AC segregation and the enrichment of H4K5ac, H4K12ac and H3K9ac on ACs depend on RbAp46/48^**LIN-53**^ **and HAT-1**

To investigate whether the corresponding histone modifying enzymes of the enriched AC PTMs facilitate *de novo* centromere formation, we performed RNA interference by injecting dsRNA of candidate histone modifier genes to L4 stage worms expressing GFP::LacI and mCherry::H2B (Figure 5A). The RNAi depletion efficiency of each gene was confirmed by live imaging (Figure S4A) or RT-qPCR (Figure S4B). Eighteen hours after dsRNA injection, L64×LacO was injected to RNAi-treated worms or untreated worms of the same stage. Embryos were dissected from injected worms, mounted for live-cell imaging to measure the AC segregation rate, and for immunofluorescence analysis to compare the CENP-A^HCP-3^ signal on ACs. Snapshots (Figure 5B) and videos (Suppl. Video3 and 4) from live-cell imaging show an example of a WT (untreated) and a *lin-53* RNAi-treated embryo that contains ACs in the first 3 consecutive cell divisions, from one-cell stage to four-cell stage. A nascent AC in a WT embryo aligned at the metaphase plate with endogenous chromosomes, attempted to segregate, but formed chromosome bridges at the first anaphase in one-to-two cell stage, and has less severe chromosome lagging during the three-to-four cell transition. For untreated controls, the percentage of one-cell embryos that have segregating ACs among all one-cell embryos with ACs is 60%, including segregating ACs with anaphase bridges. In contrast, all the nascent ACs loss their segregation competency in *lin-53* RNAi-treated embryo, and just passively remain in one of the two daughter cells during each division.

**Figure 5.**
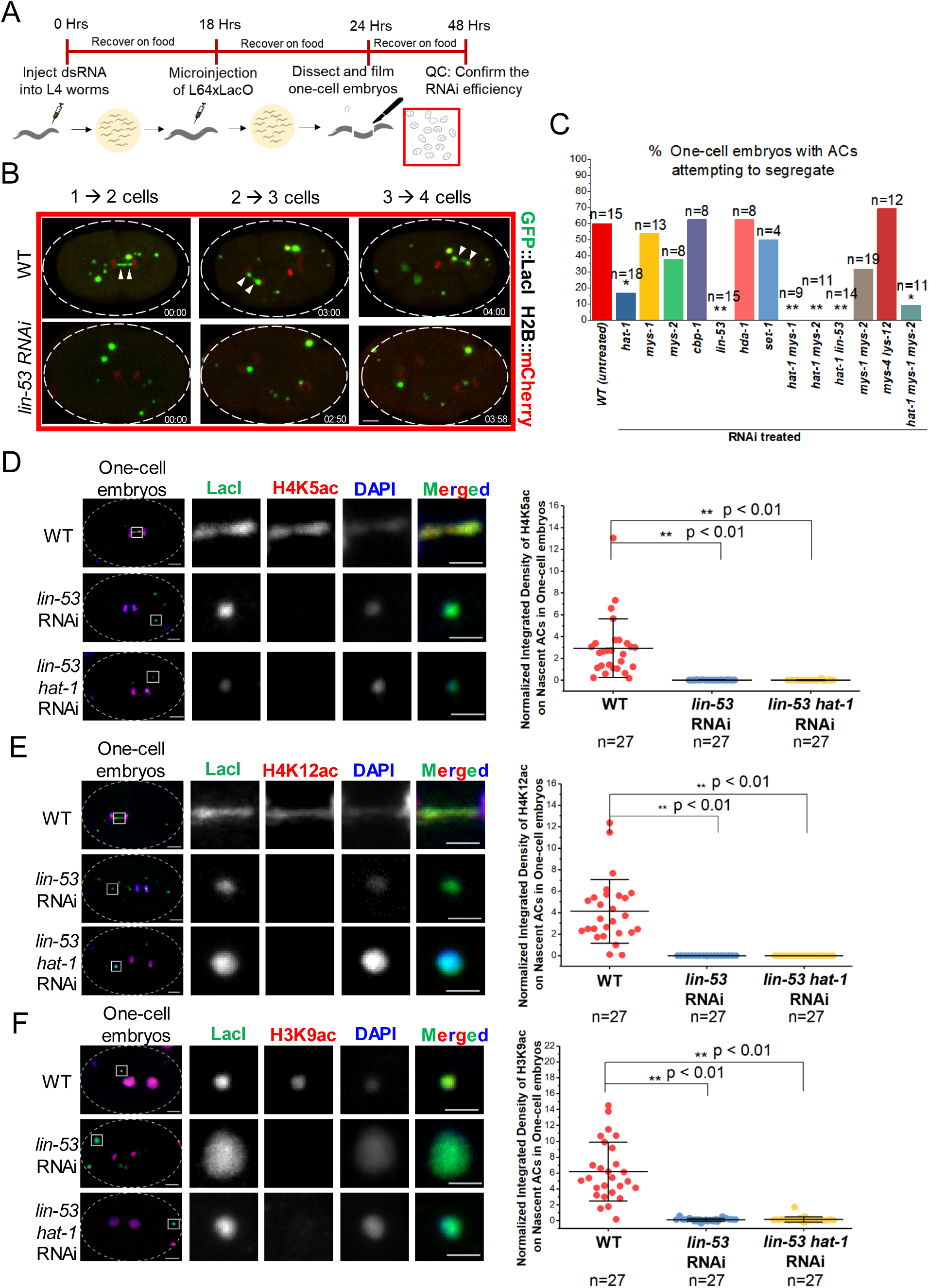
The segregation ability of nascent ACs and the enrichment of H4K5ac, H4K12ac and H3K9ac on ACs depend on RbAp46/48^LIN-53^ and HAT-1. (A) A schematic diagram of the experimental approach used to identify factors responsible for AC segregation by RNAi and live-cell imaging. (B) Representative live-cell imaging of a nascent AC that was attempting to segregate (even with anaphase bridges) in WT (untreated) and in *lin-53* RNAi-treated one-cell embryos. The time-lapse between the images was shown (mm:ss). Scale bar represents 5 μm. The same method was used for screening potential factors that affect the segregation rate of nascent ACs in one-cell embryos. (C) Quantification of AC segregation rates in WT (untreated), *hat-1, mys-1, mys-2, cbp-1, lin-53, hda-1, set-1, hat-1 mys-1* double, *hat-1 mys-2* double, *mys-1 mys-2* double, *hat-1 lin-53* double, *mys-4 lsy-12* double *and hat-1 mys-1 mys-2* triple RNAi-treated one-cell embryos. Significant differences are analyzed by the Fisher’s exact test (*, p < 0.05; **, p < 0.01). The number of samples (n) analyzed was indicated. Immunofluorescence of (D) H4K5ac, (E) H4K12ac and (F) H3K9ac on ACs in WT, *lin-53* RNAi-treated and *lin-53 hat-1* double RNAi-treated one-cell embryos. Embryos were stained with antibody against LacI (green), antibodies against a histone PTM (red) and DAPI (blue). A higher-magnification view of the ACs (white square) is shown on the right. Scale bar represents 2 μm for the magnified images. Scatter plots show the quantification of normalized integrated density of (D) H4K5ac, (E) H4K12ac and (F) H3K9ac on ACs. The integrated density of each PTM was normalized to DAPI. The number of samples (n) analyzed was indicated. The error bars represent SD. Significant differences are analyzed by student’s t-test (**, p < 0.01).

We also depleted individual histone acetyltransferases *(hat-1, cbp-1, mys-1, mys-2, lsy-12* and *mys-4*), a histone deacetylase (*hda-1*), a histone methyltransferase (*set-1*) that is responsible for H4K20me (18), or depleted them in double and triple combinations. Among all single RNAi treatments of acetyltransferases, *hat-1* RNAi significantly reduces AC segregation frequency to 17% (p < 0.05). Single RNAi of other acetyltransferases (*cbp-1, mys-1, mys-2*) does not cause any significant decrease in the AC segregation rate. Double knockdown of *hat-1 mys-1* and *hat-1 mys-2* and triple knockdown of *hat-1 mys-1 mys-2* further decrease AC segregation rates in one-cell stage embryos. These findings indicate that acetyltransferases play an essential role in the segregation of nascent ACs. MYS-1 and MYS-2 may share some overlapping acetylation targets with HAT-1, and thus have additive effects upon depletion. In contrast, the AC segregation rates in *mys-1 mys-2* knockdown and *lsy-12 mys-4* knockdown embryos, at 30% and 67%, respectively, have no significant difference with WT embryos (Figure 5C). Depletion of *hda-1* or *set-1* also did not affect AC segregation in one-cell embryos (Figure 5C).

Although RbAp46/48 and HAT-1 were conserved, and found in the same complex in many species (19-21), the physical interaction between RbAp46/48^LIN-53^ and HAT-1 has not been reported in *C. elegans* (22). We created a transgenic strain expressing GFP::HAT-1 by CRISPR-Cas9 at the endogenous locus (7). The expression of GFP::HAT-1 was observed in embryonic nuclei (Figure S4A). We also confirmed the physical interaction between RbAp46/48^LIN-53^ and HAT-1 by reciprocal co-immunoprecipitation (co-IP) using embryo extracts (Figure S5).

As RbAp46/48 is known to be a H3-H4 chaperone (23-25), we found that RbAp46/48^LIN-53^ is also essential for histone H3 deposition on nascent ACs, but surprisingly not histone H4 (Figure S4C and S4D). The levels of H4K5ac, H4K12ac and H3K9ac on nascent ACs are significantly decreased to an undetectable level in both *lin*-53 RNAi-treated and in *lin-53 hat-1* double RNAi-treated embryos (Figure 5D-F), indicating that RbAp46/48^LIN-53^ and HAT-1 may function together for depositing acetylated histones.

### HAT-1 assists RbAp46/48^LIN-53^ in *de novo* CENP-A^HCP-3^ deposition on nascent ACs

Since depletion of *hat-1, lin-53* and double depletion of *lin-53 hat-1* significantly decreased AC segregation rate, we proposed that RbAp46/48^LIN-53^ and HAT-1 are responsible for depositing CENP-A^HCP-3^-H4 pre-nucleosomes on nascent ACs after fertilization. We performed immunofluorescence analysis of CENP-A^HCP-3^ on the nascent ACs in *hat-1, lin-53* and *lin-53 hat-1* double RNAi-treated embryos. We found that the CENP-A^HCP-3^ level on ACs is significantly decreased in *hat-1* and *lin-53* RNAi-treated embryos (Figure 6A-C) and is completely abolished in *lin-53 hat-1* double RNAi-treated embryos (Figure 6A and 6D). The more severe abolishment of CENP-A^HCP-3^ in *lin-53 hat-1* double depletion suggests that HAT-1 and RbAp46/48^LIN-53^ may also function separately for recruiting CENP-A^HCP-3^, in addition to acting together in a complex.

**Figure 6.**
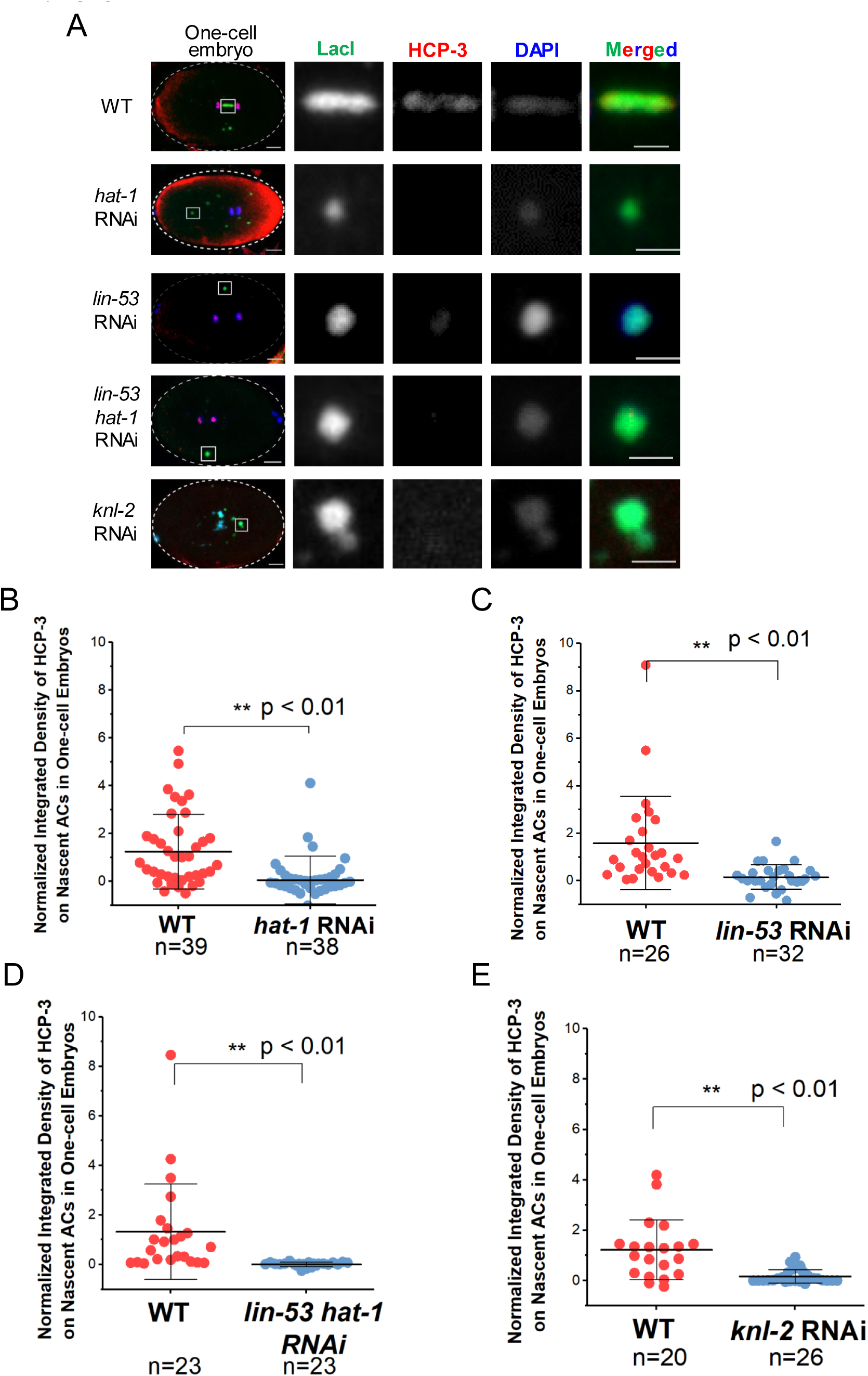
HAT-1 assists RbAp46/48^LIN-53^ in *de novo* CENP-A^HCP-3^ deposition on nascent ACs. (A) Immunofluorescence of CENP-A^HCP-3^ on ACs in WT, *hat-1* RNAi, *lin-53, lin-53 hat-1* double and *knl-2* RNAi-treated one-cell embryos. A higher-magnification view of the ACs (white square) is shown on the right. Scale bars in whole embryo images and in the magnified images represent 5 μm and 2 μm, respectively. Scatter plot shows the quantification result of the normalized integrated density of CENP-A^HCP-3^ signal on ACs in (B) *hat-1*, (C) *lin-53*, (D) *lin-53 hat-1* double and (E) *knl-2* RNAi-treated one-cell embryos, which were compared with that in WT embryos. For (B)-(E), the integrated density of CENP-A^HCP-3^ was normalized to DAPI. The number of samples (n) analyzed was indicated. The error bars represent SD. Significant differences are analyzed by the student’s t-test (**, p < 0.01).

### RbAp46/48^LIN-53^-initiated d*e novo* CENP-A^HCP-3^ deposition is required for Mis18BP1^KNL-2^ localization on ACs

Mis18BP1^KNL-2^ and CENP-A^HCP-3^ are interdependent for each other’s localization in endogenous chromosomes of *C. elegans* (26). To test if Mis18BP1^KNL-2^ is also necessary for *de novo* CENP-A^HCP-3^ deposition on nascent ACs, we depleted Mis18BP1^KNL-2^ and performed immunofluorescence analysis, which shows that *knl-2* RNAi almost completely abolished CENP-A^HCP-3^ signal on nascent ACs (Figure 6A and 6E). This indicates that Mis18BP1^KNL-2^ is also essential for CENP-A^HCP-3^ localization on both nascent ACs and endogenous centromeres. Similar to endogenous centromeres, Mis18BP1^KNL-2^ localization on nascent ACs also relies on CENP-A^HCP-3^ (Figure 7A and 7C). We simultaneously stained CENP-A^HCP-3^ and Mis18BP1^KNL-2^ on nascent ACs in one-cell embryos, and show that 62% of ACs have both CENP-A^HCP-3^ and Mis18BP1^KNL-2^, while 38% of ACs have neither of the signals. We have not found any ACs that have only CENP-A^HCP-3^ or only Mis18BP1^KNL-2^ (Figure S6A and B), which is consistent with the co-dependence of Mis18BP1^KNL-2^ and CENP-A^HCP-3^ localization (Figure 8A).

**Figure 7.**
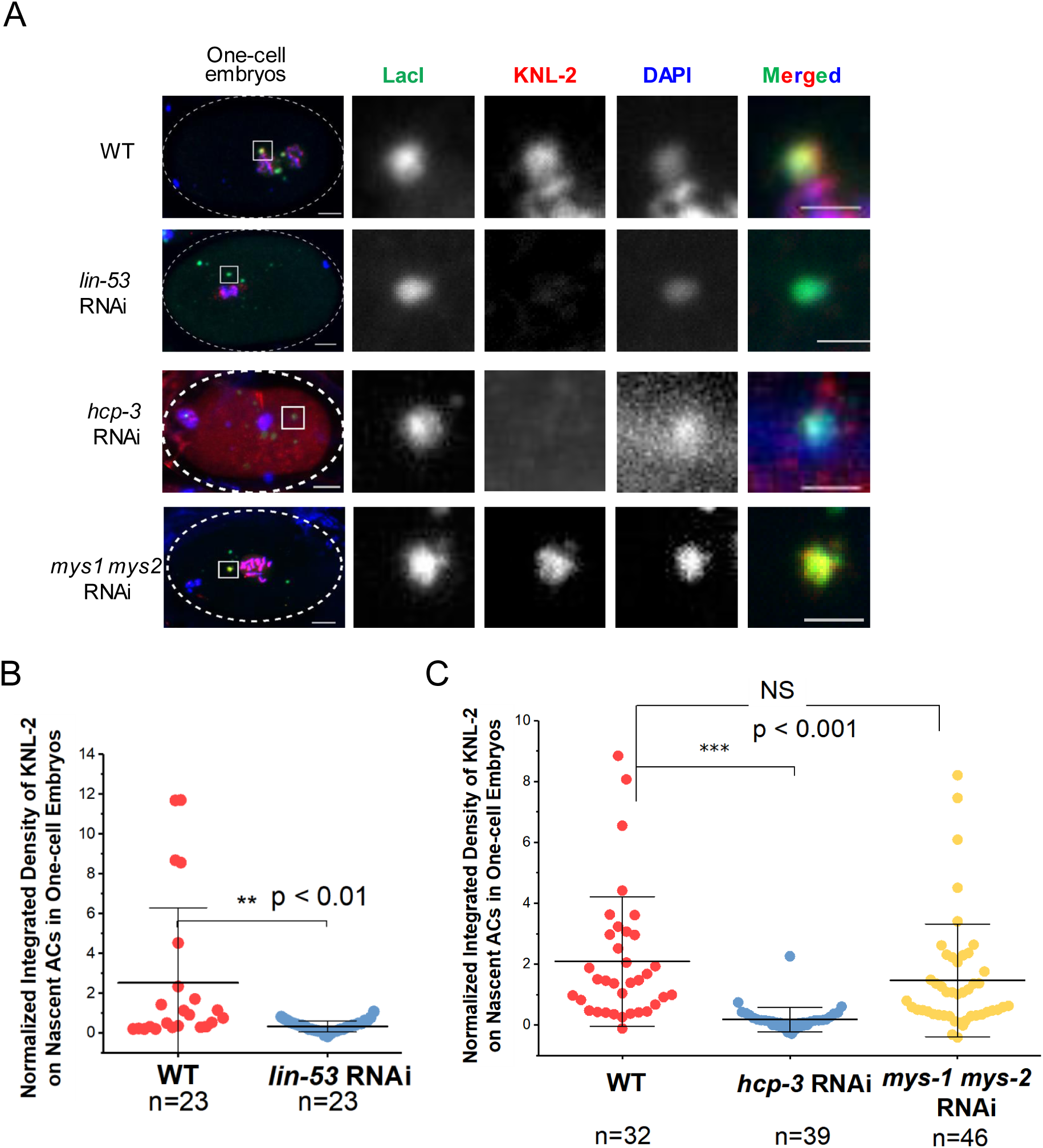
RbAp46/48^LIN-53^-initiated *de novo* CENP-A^HCP-3^ deposition is required for Mis18BP1^KNL-2^ localization. (A) Immunofluorescence of M18BP1^KNL-2^ on ACs in WT, *lin-53, hcp-3* and *mys-1 mys-2* double RNAi-treated one-cell embryos. A higher-magnification view of the ACs (white square) is shown on the right. Scale bars in whole embryo images and in the magnified images represent 5 μm and 2 μm, respectively. A scatter plot shows the quantification result of the normalized integrated density of M18BP1^KNL-2^ signal on ACs in (B) *lin-53*, (C) *hcp-3* and *mys-1 mys-2* double RNAi-treated one-cell embryos, which were compared with that in WT embryos. For (B) and (C), the integrated density of M18BP1^KNL-2^ was normalized to DAPI. The number of samples (n) analyzed was indicated. The error bars represent SD. Significant differences are analyzed by the student’s t-test (**, p<0.01; ***, p<0.001)).

**Figure 8.**
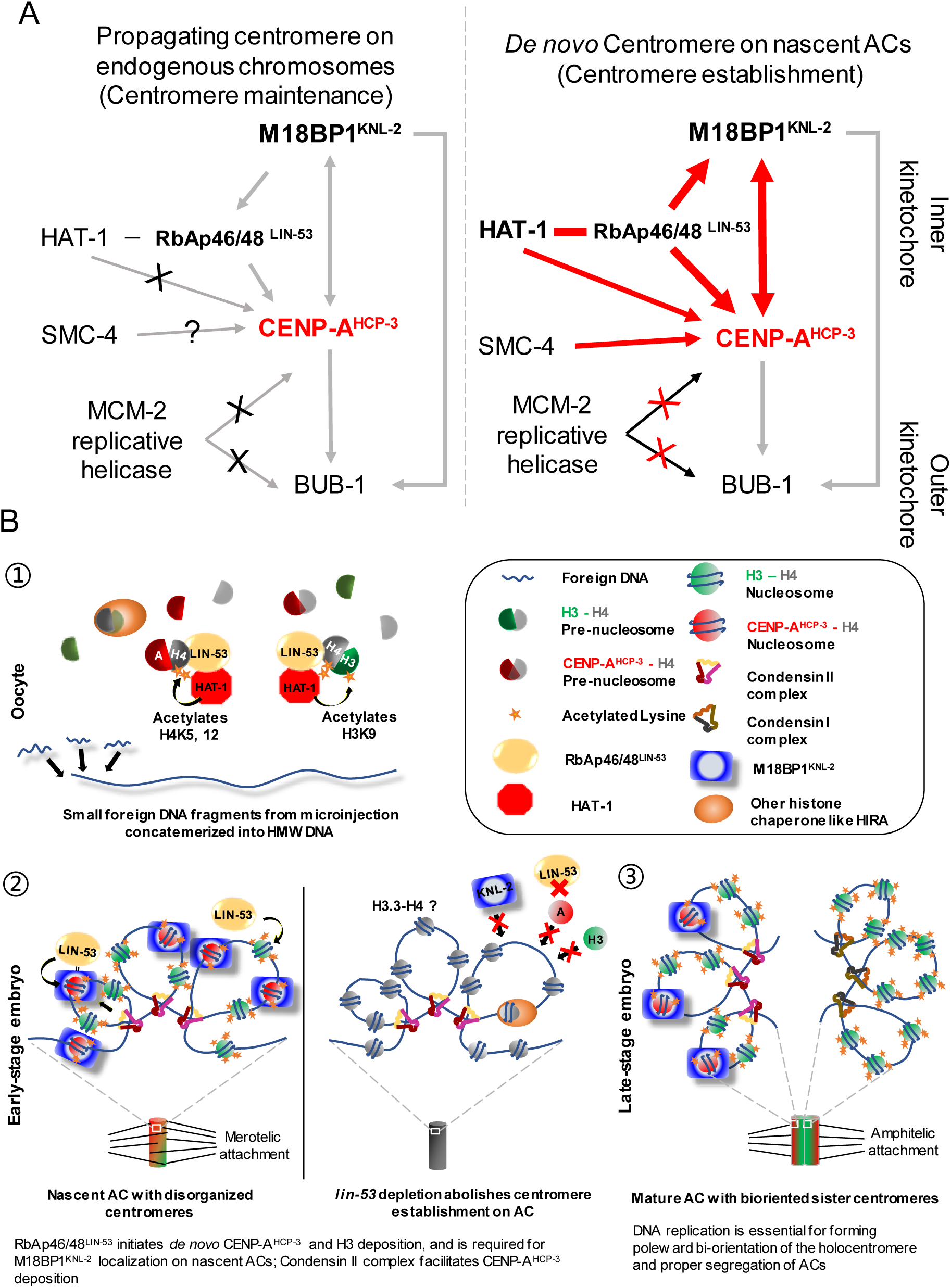
A schematic diagram of the *de novo* centromere formation in *C. elegans* embryos. (A) A schematic diagram of the centromeric localization dependency in *C. elegans* endogenous chromosomes and in *de novo* centromere formation on ACs. A→B means B’s localization is dependent on A, but not the other way around. Gray arrows indicate findings from other studies. Red arrows indicate the findings from this study. Bold arrows indicate that the effects are very severe. The line between two factors indicates that they have physical interaction. (B) The proposed process of artificial chromosome formation in *C. elegans* gonad. 1. Firstly, small foreign DNA fragments from microinjection concatemerizes into HMW DNA arrays in the oocyte cells. RbAp46/48^LIN-53^-HAT-1 complex acetylates H3-H4 and CENP-A-H4 pre-nucleosomes at H4K5, H4K12 and H3K9, which contributes to the hyperacetylation of nascent ACs. 2. Secondly, RbAp46/48^LIN-53^ initiates *de novo* CENP-A^HCP-3^ and H3 deposition, and RbAp46/48^LIN-53^ is required for M18BP1^KNL-2^ localization on HMW DNA; Condensin II complex also facilitates CENP-A^HCP-3^ deposition. Chromatinization and centromerization of the HMW DNA generates nascent ACs. Nascent ACs have DNA replication defects and lack bi-oriented sister kinetochores, which could lead to merotelic attachments to the mitotic spindle and chromosome bridging (in early embryonic cells). 3. Finally, DNA replication efficiency gradually improves on ACs, and ACs “mature” by late embryonic cell stage. In matured ACs, bi-oriented sister kinetochores allow amphitelic attachment of spindles and proper segregation.

However, on endogenous chromosomes, *lin-53* RNAi only reduced CENP-A^HCP-3^ level but did not affect Mis18BP1^KNL-2^ level (27). We monitored the M18BP1^KNL-2^ signal on nascent ACs in *lin-53* RNAi-treated embryos. Surprisingly, at *de novo* centromeres on nascent ACs, RbAp46/48^LIN-53^ depletion also leads to the loss of initial Mis18BP1^KNL-2^ deposition (Figure 7A and 7B), which suggests that while Mis18BP1^KNL-2^ could be a self-directing factor for centromere maintenance in the existing centromeres, it is downstream of RbAp46/48^LIN-53^ in *de novo* centromere establishment (Figure 8A).

In human cells, MYST2 has been described as an interactor with Mis18 complex for regulating CENP-A deposition (28). However, the function of MYS family proteins on centromere chromatin has not been reported previously in *C. elegans*. Since we found that MYS-1 and MYS-2 can partially complement HAT-1 in facilitating AC segregation, we tested whether double knockdown of *mys-1* and *mys-2* prevents Mis18BP1^KNL-2^ localization on nascent ACs. However, the Mis18BP1^KNL-2^ signal on nascent ACs in *mys-1* and *mys-2* double depleted embryos shows no significant difference as compared with WT (untreated) embryos (Figure 7A and 7C).

## DISCUSSION

In our previous study, we observed that foreign circular, supercoiled plasmid DNA injected into *C. elegans* gonad forms ACs in embryonic cells after 4-8 hours of microinjection (6). ACs gradually acquire segregation competency after they go through several cell divisions (6). Since then, we have tested AC formation by injecting different DNA forms, including linearized plasmid DNA and linearized plasmid DNA mixed with sheared salmon or enzyme-digested yeast genomic DNA (Lin and Yuen, submitted back-to-back). We found that it is more efficient to form larger ACs by concatemerization from linear DNA than from circular DNA, based on the foci size of GFP::LacI, which binds to the injected LacO arrays, in the embryos (Figures S1D and S1E). This may suggest that it is more efficient to fuse linear foreign DNA fragments by the non-homologous end joining (NHEJ) pathway than fusing circular DNA (4). We also found that ACs generated by injecting linear DNA (L64×LacO) acquires segregation ability significantly faster than those generated by injecting circular DNA (Figure S1F), possibly due to the larger ACs it produces. By injecting a complex DNA mixture from sheared salmon sperm DNA without the LacO repeat sequence, we confirmed that *de novo* CENP-A^HCP-3^ deposition can also occur on “complex” AC without LacO sequences (Figure S1G and S1H). However, the AC segregation rates in repetitive ACs and complex ACs have no significant difference (Lin and Yuen, submitted back-to-back).

Chromatinized ACs were formed in fertilized embryos a few hours after microinjection of foreign circular DNA (6) and linear DNA. Histone H2B and centromeric protein CENP-A^HCP-3^ signals on ACs were detectable in fertilized embryos (Figure 1 B-C). This is consistent with the finding that major sperm proteins trigger nuclear membrane breakdown (29) and release the nuclear-localized histones and centromeric proteins to the cytoplasm to allow chromatinization and centromerization on the HMW-DNA arrays, which are initially located in the cytoplasm. The whole process of AC formation has been summarized in Figure 8B.

Interestingly, in wild-type, the level of CENP-A^HCP-3^ (normalized to the amount of DNA based on DAPI staining) on nascent ACs formed from linear plasmid DNA is comparable to that on endogenous chromosomes in one-cell embryos (Figure S2K). This observation is slightly different from the nascent ACs formed from circular DNA, in which the level of CENP-A^HCP-3^ on ACs in one-cell stage is lower than that on endogenous chromosomes, whilst CENP-A^HCP-3^ signal on ACs increases quickly in the first few cell cycles to become comparable with that in endogenous chromosomes in 17-32 cell stage (17). This difference could be due to the difference in size or structure of the ACs (Figure S1D-F).

DNA replication is required for chromosome condensation during prophase (the first cell cycle) and chromosome decondensation during anaphase (the second cell cycle) in *C. elegans* embryos (9). HU-treated embryos have been described to cause excessive chromatin bridge formation (30), with persistent association with condensin II subunit SMC-4 (Figure S2A and S2B), which resembles the phenomenon of nascent AC segregation with bridges. We examined the DNA replication status on nascent ACs in one-cell stage and in later stage embryos. The efficiency of incorporating EdU on the newly formed ACs is only about 32% of that of the endogenous chromosomes in one-cell embryos (Figure 2C and D), suggesting that DNA replication is less efficient on ACs. We demonstrated that MCM-4::mCherry is still associated with ACs at the first metaphase, which further indicates that the replication of AC had not completed when the cells entered first mitosis (Figure S2C and S2D). We have ruled out the possibility that LacI::GFP-tethering on ACs restrains replication initiation or replication fork movement, as the lagging AC is still observed in cells without LacI::GFP expression (Figure S2E). In *C. elegans*, DNA replication origins contain H3K4me2 enrichment (31), whereas H3K4me2 is absent on nascent ACs. The underlying sequences and chromatin environment of ACs could be the reason for the less efficient DNA replication on nascent ACs.

During DNA replication, nucleosomes are disassembled from the parental DNA strand ahead of the DNA replication machinery (32). MCM-2 has been recently proposed to be able to chaperone H3-H4 and CENP-A-H4 dimers, for replenishing them to the sister chromatins behind the replication forks (11). However, *mcm-2* RNAi did not reduce the level of *de novo* CENP-A^HCP-3^, normalized to DNA (DAPI), on nascent ACs (Figure 2E and 2F), and their ability to recruit outer kinetochore and spindle checkpoint component BUB-1 (Figure S2J). Thus, we postulate that *de novo* centromere formation *per se* is independent of DNA replication. This is similar to the case in human cells, where CENP-A is duplicated on chromatin before and independent of DNA replication (33). Although it is not clear when new CENP-A^HCP-3^ is loaded on the holocentromere during the cell cycle in *C. elegans*, we speculate that the CENP-A^HCP-3^ level in *C. elegans* correlates with the amount of DNA, as observed in previous study (34), and from the comparison in wild-type and *mcm-2* RNAi-treated embryos (Figure 2E and 2F). On the other hand, CENP-A^cse4^ in budding yeast is turned over and reloaded to sister chromatids in S phase, dependent on DNA replication (35,36).

Condensin II, but not condensin I, is specifically enriched at the centromeres and has been found to promote CENP-A deposition in human cells and *Xenopus* oocyte extracts (15,16,37). In human cells, the interaction between condensin II and HJURP is needed for HJURP’s centromeric localization, and for depositing new CENP-A (15). In contrast, depleting condensin II in *Xenopus* oocyte extracts reduces the CENP-A level at centromere, but not the HJURP level. In *C. elegans*, condensin II subunits also co-localize with CENP-A^HCP-3^ starting at prometaphase in embryonic cells (13). We found that depleting condensin II component SMC-4 reduces *de novo* CENP-A^HCP-3^ deposition on newly formed ACs, indicating that condensin II subunit SMC-4 facilitates *de novo* centromere formation. However, we could not detect physical interaction between condensin II subunit SMC-4 and CENP-A^HCP-3^ chaperone RbAp46/48^LIN-53^ (Figure S5A).

We screened for candidate histone PTMs on newly formed ACs that coincide with *de novo* centromere formation to elucidate the cellular pathways involved, because histone PTMs can signal downstream histone deposition, gene expression and chromatin condensation. H4K20me is one of the enriched PTM on nascent ACs. In human and chicken DT-40 cells, H4K20me was reported as a histone PTM on CENP-A nucleosomes enriched at centromeres, which is essential for CENP-T localization (38). In *C. elegans*, H4K20 is monomethylated by methyltransferase SET-1(18). However, the AC segregation rate in *set-1* RNAi-treated embryos shows no significant difference from that in untreated embryos (Figure 5C), indicating that H4K20me could be a potential CENP-A^HCP-3^ post-deposited marker, which may not be essential for *de novo* CENP-A^HCP-3^ deposition.

We found that histone acetylation on H3K9, H4K5 and H4K12 was significantly enriched on nascent ACs formed from linear DNA, which is consistent with the previously found acetylated H3K9 and H4 (on K5, 8, 12, or 16) on ACs from circular injected DNA (17). H3K9ac has been reported to be associated with the deposition of newly synthesized histones H3 in *Tetrahymena*, while the involved acetyltransferase is not clear (39). In human cells, H3K9ac induces *de novo* CENP-A deposition on alphoid DNA at ectopic site and is compatible with centromere functioning (40). Notably, in species as divergent as humans, *Drosophila, Tetrahymena* and yeast, newly synthesized and newly deposited H4 are acetylated in a conserved pattern at lysines 5 and 12 before its association with DNA (20,39,41). Pre-nucleosomal H4 is di-acetylated at K5 and K12 by HAT-1 in human cells, *DT-40* cells and yeast, through forming a complex with H4 chaperone, RbAp46/48 (21,23,42). in fruit fly cells (43), and is essential for the viability and genome stability in mice (44). Double knockdown of *hat-1 mys-1* or *hat-1 mys-2* abolishes AC segregation during the first mitosis in *C. elegans, while mys-1 mys-2* double depletion causes an insignificant reduction of the AC segregation rate (Figure 5C). Consistently, MYS-1 MYS-2 double depletion does not reduce the level of M18BP1^KNL-2^ on nascent ACs (Figure 7A and 7C). Both *mys-1* RNAi and *mys-2* RNAi enhance the *hat-1* RNAi effect on AC missegregation, suggesting that MYS-1 and MYS-2 might have overlapping targets with HAT-1.

*lin-53* depletion also significantly reduces histone H3 level on nascent ACs, but surprisingly, did not affect the level of histone H4 (Figure S4D). The timing suggests that CENP-A^HCP-3^ deposition may occur at the same time as canonical nucleosome assembly, and RbAp46/48^LIN-53^ is required for both CENP-A^HCP-3^ and H3 assembly. We propose that RbAp46/48^LIN-53^ may load acetylated nucleosomes to nascent chromatin, while non-acetylated histone H4, which could form a tetramer with histone H3.3, could be deposited to nascent ACs through other histone chaperones, like HIRA (45,46). Indeed, a previous study has shown that human Hat1-RbAp46 complex binds and acetylates H4 in H3.1-H4 complex more efficiently than that in H3.3-H4 complex (47). Hence, the depletion of lin-53 leads to a significant decrease in H4K5ac and H4K12ac level (Figure 5D and 5E), but the total H4 level remains unchanged (Figure S4D). A higher proportion of unacetylated H4 is on nascent ACs, which is consistent with the idea that RbAp46/48^LIN-53^ preferentially deposits acetylated H4 on AC in *C. elegans*.

We observed a significant reduction of the AC segregation rate in *hat-1* RNAi-, *lin-53* RNAi- *and lin-53 hat-1* double RNAi*-*treated *embryos*, indicating that RbAp46/48^LIN-53^ and HAT-1 are both involved in *de novo* centromere formation (Figure 5C). In chicken DT-40 cells, RbAp48 is essential for new CENP-A deposition to the centromere, which cooperates with HAT-1 to acetylate pre-nucleosomal CENP-A-H4 complex at H4K5 and K12 (48). In *Drosophila*, an *in vitro* experiment shows that RbAp48 alone is sufficient to assemble CENP-A^CID^-H4 to the naked DNA (49). New evidence shows that HAT-1 interacts directly with CENP-A^CID^ in *Drosophila*. The depletion of HAT-1 reduces the efficiency of new CENP-A^CID^ deposition significantly (43). Nevertheless, in fission yeast, RbAp46/48^Mis16^ and HJURP^SCM3^ are present in the same complex, where RbAp46/48^Mis16^ distinguishes CENP-A-H4 from H3-H4 by recognizing HJURP^SCM3^ and H4 independently (50). In HeLa cells, RNAi knockdown of RbAp46/48 reduces ectopic loading of CENP-A^S68E^ mutant on the chromosome arms, which cannot bind to HJURP, suggesting that RbAp46/48 may deposit CENP-A^S68E^-H4 to ectopic loci in the absence of HJURP (51). However, in oocyte extracts of *Xenopus*, RbAp48 depletion does not affect CENP-A incorporation to the centromere on the sperm chromatin but causes ectopic CENP-A deposition (16,48). In *C. elegans* ACs, we found that both RbAp46/48^LIN-53^ and HAT-1 are critical for *de novo* CENP-A^HCP-3^ deposition. RbAp46/48^LIN-53^ may deposit CENP-A^HCP-3^ onto foreign DNA that has not been fully occupied by H3 or H3.3 nucleosomes (52). The enriched RNA polymerase II on the nascent ACs (17) might further create an open chromatin environment and generate nucleosome gaps that preferentially accumulate CENP-A^HCP-3^ through RbAp46/48^LIN-53^ loading (53).

M18BP1^KNL-2^, conserved in *C. elegans*, is upstream of RbAp46/48^LIN-53^, and both are essential for CENP-A^HCP-3^ deposition in endogenous centromeres (26,27). In fission yeast, Mis-18, RbAp46/48^Mis16^ and HJURP^Scm3^ have been found in the same complex for depositing CENP-A^Cnp1^ to the centromere (50,54,55). In humans and vertebrates, CENP-A-HJURP relies on the MIS18 complex (including MIS-18α, MIS-18β and M18BP1) for centromeric targeting (56,57). Tethering LacI-fused M18BP1 or Mis-18β to the LacO region promotes new CENP-A deposition by recruiting HJURP to the tethered locus (56), similar to tethering HJURP to an ectopic locus (58). M18BP1^KNL-2^, as a priming factor, is anticipated to anchor to the existing centromeres for directing new CENP-A loading. Since *C. elegans* has no HJURP, Mis18α nor Mis18β, M18BP1^KNL-2^ alone is possibly sufficient to direct RbAp46/48^LIN-53^ for depositing CENP-A^HCP-3^ to existing centromeres. Here, we show that depleting M18BP1^KNL-2^ also significantly reduced CENP-A^HCP-3^ level on nascent ACs (Figure 5A and 5G), consistent with its effect on endogenous chromosomes. The inter-dependency between M18BP1^KNL-2^ and CENP-A is also true for both endogenous chromosomes (26,27) and on nascent ACs. In RbAp46/48^LIN-53^-depleted embryos, M18BP1^KNL-2^ was not able to localize to the nascent ACs without any pre-seeded CENP-A^HCP-3^ (Figure 5A, 5G and Figure S6A and S6B). This finding strongly indicates that RbAp46/48^LIN-53^ initiates CENP-A^HCP-3^ nucleosome assembly on foreign DNA, which lays the foundation for loading of other kinetochore proteins. The centromeric localization dependency in *C. elegans* endogenous chromosomes and in *de novo* centromere formation on ACs has been summarized and compared (Figure 8A). It will be of great interest to know if RbAp46/48^LIN-53^ and M18BP1^KNL-2^ form a complex to deposit pre-nucleosomal CENP-A-H4, in which RbAp46/48^LIN-53^ is released from the centromere after the CENP-A^HCP-3^ deposition, while M18BP1^KNL-2^ is retained on the chromatin for CENP-A^HCP-3^ stabilization and for recruiting outer kinetochore proteins.

Phosphorylation of CENP-A at Ser68 has been proposed to be important for preventing premature HJURP binding at metaphase in HeLa cells (51). The phospho-mimicking mutant, CENP-A^S68Q^, reduces its affinity to HJURP (51,59), but it is still able to support centromere function and long-term centromere maintenance in human RPE-1 cells (59). However, mutating the Ser68 residue to alanine, as a phospho-dead mutant, causes continuous CENP-A binding to HJURP and ectopic CENP-A deposition (51). Interestingly, this site is evolutionarily conserved in most eukaryotes except in *C. elegans* and budding yeast, where these CENP-A homologues have an alanine at this position. Hence, budding yeast HJURP^Scm3^ constitutively binds to CENP-A^Cse4^ at centromeres throughout the cell cycle in budding yeast (35). Therefore, in budding yeast and *C. elegans*, this serine to alanine mutation may allow HJURP and RbAp46/48 to be functionally redundant in persistent binding to the CENP-A/H4 dimer. In budding yeast, CENP-A^cse4^ propagation relies on HJURP^Scm3^, and as a result, RbAp46/48^HAT2p^ can be a non-essential protein in this species (60). Similarly, *C. elegans* can afford losing HJURP, as CENP-A^HCP-3^ propagation depends on RbAp46/48^LIN-53^ instead. RbAp46/48^LIN-53^ does not rely on pre-existing CENP-A in order to deposit CENP-A^HCP-3^ on foreign DNA. Furthermore, budding yeast and *C. elegans* have similar CENP-A propagation mechanisms, which are different from CENP-A propagation in human cells. For instance, pre-existing centromeric CENP-A^Cse4^ is completely turned over in S phase in budding yeast. While the cell cycle stage of CENP-A^HCP-3^ turnover is unknown in *C. elegans*, it is also almost completely turned over (34,35). In yeast and worms, they tend to maintain a consistent amount of CENP-A per chromatin or total DNA. Potentially, budding yeast and *C. elegans* have developed an alternative pathway of regulating CENP-A propagation, in which the cue for new CENP-A deposition is not only dependent on pre-existing CENP-A. This, combined with promiscuous CENP-A^HCP-3^ deposition facilitated by RbAp46/48^LIN-53^, allows ACs to be formed easily in *C. elegans*, which provides a convenient model for the study of *de novo* centromere formation.

## Supporting information

Supplementary table&figure egends

## ACKNOWLEDGEMENT

We thank J Dumont, KM Chan, R Ng, Y Zhai, and W Den for critical reading of earlier versions of the manuscript, and the Yuen lab for discussion.

## REFERENCES

1. Marshall, O.J., Chueh, A.C., Wong, L.H. and Choo, K.H. (2008) Neocentromeres: new insights into centromere structure, disease development, and karyotype evolution. American journal of human genetics, 82, 261–282.

2. Catania, S., Pidoux, A.L. and Allshire, R.C. (2015) Sequence features and transcriptional stalling within centromere DNA promote establishment of CENP-A chromatin. PLoS genetics, 11, e1004986.

3. Harrington, J.J., Van Bokkelen, G., Mays, R.W., Gustashaw, K. and Willard, H.F. (1997) Formation of de novo centromeres and construction of first-generation human artificial microchromosomes. Nature genetics, 15, 345–355.

4. Stinchcomb, D.T., Shaw, J.E., Carr, S.H. and Hirsh, D. (1985) Extrachromosomal DNA transformation of Caenorhabditis elegans. Molecular and cellular biology, 5, 3484–3496.

5. Mello, C.C., Kramer, J.M., Stinchcomb, D. and Ambros, V. (1991) Efficient gene transfer in C.elegans: extrachromosomal maintenance and integration of transforming sequences. The EMBO journal, 10, 3959–3970.

6. Yuen, K.W., Nabeshima, K., Oegema, K. and Desai, A. (2011) Rapid de novo centromere formation occurs independently of heterochromatin protein 1 in C. elegans embryos. Current biology: CB, 21, 1800–1807.

7. Dickinson, D.J., Pani, A.M., Heppert, J.K., Higgins, C.D. and Goldstein, B. (2015) Streamlined Genome Engineering with a Self-Excising Drug Selection Cassette. Genetics.

8. Carvalho, A., Olson, S.K., Gutierrez, E., Zhang, K., Noble, L.B., Zanin, E., Desai, A., Groisman, A. and Oegema, K. (2011) Acute drug treatment in the early C. elegans embryo. PloS one, 6, e24656.

9. Sonneville, R., Craig, G., Labib, K., Gartner, A. and Blow, J.J. (2015) Both Chromosome Decondensation and Condensation Are Dependent on DNA Replication in C. elegans Embryos. Cell reports, 12, 405–417.

10. Sonneville, R., Querenet, M., Craig, A., Gartner, A. and Blow, J.J. (2012) The dynamics of replication licensing in live Caenorhabditis elegans embryos. The Journal of cell biology, 196, 233–246.

11. Huang, H., Stromme, C.B., Saredi, G., Hodl, M., Strandsby, A., Gonzalez-Aguilera, C., Chen, S., Groth, A. and Patel, D.J. (2015) A unique binding mode enables MCM2 to chaperone histones H3-H4 at replication forks. Nature structural & molecular biology, 22, 618–626.

12. Kaitna, S., Pasierbek, P., Jantsch, M., Loidl, J. and Glotzer, M. (2002) The aurora B kinase AIR-2 regulates kinetochores during mitosis and is required for separation of homologous chromosomes during meiosis. Current Biology, 12, 798–812.

13. Csankovszki, G., Collette, K., Spahl, K., Carey, J., Snyder, M., Petty, E., Patel, U., Tabuchi, T., Liu, H., McLeod, I. et al. (2009) Three distinct condensin complexes control C. elegans chromosome dynamics. Current biology: CB, 19, 9–19.

14. Hagstrom, K.A., Holmes, V.F., Cozzarelli, N.R. and Meyer, B.J. (2002) C. elegans condensin promotes mitotic chromosome architecture, centromere organization, and sister chromatid segregation during mitosis and meiosis. Genes & development, 16, 729–742.

15. Barnhart-Dailey, M.C., Trivedi, P., Stukenberg, P.T. and Foltz, D.R. (2017) HJURP interaction with the condensin II complex during G1 promotes CENP-A deposition. Molecular biology of the cell, 28, 54–64.

16. Bernad, R., Sanchez, P., Rivera, T., Rodriguez-Corsino, M., Boyarchuk, E., Vassias, I., Ray-Gallet, D., Arnaoutov, A., Dasso, M., Almouzni, G. et al. (2011) Xenopus HJURP and condensin II are required for CENP-A assembly. The Journal of cell biology, 192, 569–582.

17. Zhu, J., Cheng, K.C.L. and Yuen, K.W.Y. (2018) Histone H3K9 and H4 Acetylations and Transcription Facilitate the Initial CENP-A(HCP-3) Deposition and De Novo Centromere Establishment in Caenorhabditis elegans Artificial Chromosomes. Epigenetics & chromatin, 11, 16.

18. Vielle, A., Lang, J., Dong, Y., Ercan, S., Kotwaliwale, C., Rechtsteiner, A., Appert, A., Chen, Q.B., Dose, A., Egelhofer, T. et al. (2012) H4K20me1 contributes to downregulation of X-linked genes for C. elegans dosage compensation. PLoS genetics, 8, e1002933.

19. Tong, K., Keller, T., Hoffman, C.S. and Annunziato, A.T. (2012) Schizosaccharomyces pombe Hat1 (Kat1) is associated with Mis16 and is required for telomeric silencing. Eukaryotic cell, 11, 1095–1103.

20. Ruiz-Garcia, A.B., Sendra, R., Galiana, M., Pamblanco, M., Perez-Ortin, J.E. and Tordera, V. (1998) HAT1 and HAT2 proteins are components of a yeast nuclear histone acetyltransferase enzyme specific for free histone H4. The Journal of biological chemistry, 273, 12599–12605.

21. Verreault, A., Kaufman, P.D., Kobayashi, R. and Stillman, B. (1998) Nucleosomal DNA regulates the core-histone-binding subunit of the human Hat1 acetyltransferase. Current biology: CB, 8, 96–108.

22. Lu, X.W. and Horvitz, H.R. (1998) lin-35 and lin-53, two genes that antagonize a C-elegans Ras pathway, encode proteins similar to Rb and its binding protein RbAp48. Cell, 95, 981–991.

23. Li, Y., Zhang, L., Liu, T., Chai, C., Fang, Q., Wu, H., Agudelo Garcia, P.A., Han, Z., Zong, S., Yu, Y. et al. (2014) Hat2p recognizes the histone H3 tail to specify the acetylation of the newly synthesized H3/H4 heterodimer by the Hat1p/Hat2p complex. Genes & development, 28, 1217–1227.

24. Satrimafitrah, P., Barman, H.K., Ahmad, A., Nishitoh, H., Nakayama, T., Fukagawa, T. and Takami, Y. (2016) RbAp48 is essential for viability of vertebrate cells and plays a role in chromosome stability. Chromosome research: an international journal on the molecular, supramolecular and evolutionary aspects of chromosome biology, 24, 161–173.

25. Campos, E.I., Fillingham, J., Li, G., Zheng, H., Voigt, P., Kuo, W.H., Seepany, H., Gao, Z., Day, L.A., Greenblatt, J.F. et al. (2010) The program for processing newly synthesized histones H3.1 and H4. Nature structural & molecular biology, 17, 1343–1351.

26. Maddox, P.S., Hyndman, F., Monen, J., Oegema, K. and Desai, A. (2007) Functional genomics identifies a Myb domain-containing protein family required for assembly of CENP-A chromatin. The Journal of cell biology, 176, 757–763.

27. Lee, B.C., Lin, Z. and Yuen, K.W. (2016) RbAp46/48(LIN-53) Is Required for Holocentromere Assembly in Caenorhabditis elegans. Cell reports, 14, 1819–1828.

28. Ohzeki, J., Shono, N., Otake, K., Martins, N.M., Kugou, K., Kimura, H., Nagase, T., Larionov, V., Earnshaw, W.C. and Masumoto, H. (2016) KAT7/HBO1/MYST2 Regulates CENP-A Chromatin Assembly by Antagonizing Suv39h1-Mediated Centromere Inactivation. Developmental cell, 37, 413–427.

29. Han, S.M., Cottee, P.A. and Miller, M.A. (2010) Sperm and oocyte communication mechanisms controlling C. elegans fertility. Dev Dyn, 239, 1265–1281.

30. Brauchle, M., Baumer, K. and Gonczy, P. (2003) Differential activation of the DNA replication checkpoint contributes to asynchrony of cell division in C. elegans embryos. Current biology: CB, 13, 819–827.

31. Pourkarimi, E., Bellush, J.M. and Whitehouse, I. (2016) Spatiotemporal coupling and decoupling of gene transcription with DNA replication origins during embryogenesis in C. elegans. Elife, 5.

32. Vijayraghavan, S. and Schwacha, A. (2012) The eukaryotic Mcm2-7 replicative helicase. Subcell Biochem, 62, 113–134.

33. Bui, M., Dimitriadis, E.K., Hoischen, C., An, E., Quenet, D., Giebe, S., Nita-Lazar, A., Diekmann, S. and Dalal, Y. (2012) Cell-cycle-dependent structural transitions in the human CENP-A nucleosome in vivo. Cell, 150, 317–326.

34. Gassmann, R., Rechtsteiner, A., Yuen, K.W., Muroyama, A., Egelhofer, T., Gaydos, L., Barron, F., Maddox, P., Essex, A., Monen, J. et al. (2012) An inverse relationship to germline transcription defines centromeric chromatin in C. elegans. Nature, 484, 534–537.

35. Wisniewski, J., Hajj, B., Chen, J.J., Mizuguchi, G., Xiao, H., Wei, D., Dahan, M. and Wu, C. (2014) Imaging the fate of histone Cse4 reveals de novo replacement in S phase and subsequent stable residence at centromeres. Elife, 3.

36. Pearson, C.G., Yeh, E., Gardner, M., Odde, D., Salmon, E.D. and Bloom, K. (2004) Stable Kinetochore-Microtubule Attachment Constrains Centromere Positioning in Metaphase. 14, 1962–1967.

37. Ono, T., Fang, Y., Spector, D.L. and Hirano, T. (2004) Spatial and temporal regulation of Condensins I and II in mitotic chromosome assembly in human cells. Molecular biology of the cell, 15, 3296–3308.

38. Hori, T., Shang, W.H., Toyoda, A., Misu, S., Monma, N., Ikeo, K., Molina, O., Vargiu, G., Fujiyama, A., Kimura, H. et al. (2014) Histone H4 Lys 20 monomethylation of the CENP-A nucleosome is essential for kinetochore assembly. Developmental cell, 29, 740–749.

39. Sobel, R.E., Cook, R.G., Perry, C.A., Annunziato, A.T. and Allis, C.D. (1995) Conservation of deposition-related acetylation sites in newly synthesized histones H3 and H4. Proceedings of the National Academy of Sciences of the United States of America, 92, 1237–1241.

40. Ohzeki, J., Bergmann, J.H., Kouprina, N., Noskov, V.N., Nakano, M., Kimura, H., Earnshaw, W.C., Larionov, V. and Masumoto, H. (2012) Breaking the HAC Barrier: histone H3K9 acetyl/methyl balance regulates CENP-A assembly. The EMBO journal, 31, 2391–2402.

41. Ejlassi-Lassallette, A., Mocquard, E., Arnaud, M.C. and Thiriet, C. (2011) H4 replication-dependent diacetylation and Hat1 promote S-phase chromatin assembly in vivo. Molecular biology of the cell, 22, 245–255.

42. Murzina, N.V., Pei, X.Y., Zhang, W., Sparkes, M., Vicente-Garcia, J., Pratap, J.V., McLaughlin, S.H., Ben-Shahar, T.R., Verreault, A., Luisi, B.F. et al. (2008) Structural basis for the recognition of histone H4 by the histone-chaperone RbAp46. Structure, 16, 1077–1085.

43. Boltengagen, M., Huang, A., Boltengagen, A., Trixl, L., Lindner, H., Kremser, L., Offterdinger, M. and Lusser, A. (2015) A novel role for the histone acetyltransferase Hat1 in the CENP-A/CID assembly pathway in Drosophila melanogaster. Nucleic acids research.

44. Nagarajan, P., Ge, Z., Sirbu, B., Doughty, C., Agudelo Garcia, P.A., Schlederer, M., Annunziato, A.T., Cortez, D., Kenner, L. and Parthun, M.R. (2013) Histone acetyl transferase 1 is essential for mammalian development, genome stability, and the processing of newly synthesized histones H3 and H4. PLoS genetics, 9, e1003518.

45. Tagami, H., Ray-Gallet, D., Almouzni, G. and Nakatani, Y. (2004) Histone H3.1 and H3.3 Complexes Mediate Nucleosome Assembly Pathways Dependent or Independent of DNA Synthesis. Cell, 116, 51–61.

46. Ricketts, M.D., Dasgupta, N., Fan, J., Han, J., Gerace, M., Tang, Y., Black, B.E., Adams, P.D. and Marmorstein, R. (2019) The HIRA histone chaperone complex subunit UBN1 harbors H3/H4- and DNA-binding activity. The Journal of biological chemistry, 294, 9239–9259.

47. Zhang, H., Han, J., Kang, B., Burgess, R. and Zhang, Z. (2012) Human Histone Acetyltransferase 1 Protein Preferentially Acetylates H4 Histone Molecules in H3.1-H4 over H3.3-H4. 287, 6573–6581.

48. Shang, W.H., Hori, T., Westhorpe, F.G., Godek, K.M., Toyoda, A., Misu, S., Monma, N., Ikeo, K., Carroll, C.W., Takami, Y. et al. (2016) Acetylation of histone H4 lysine 5 and 12 is required for CENP-A deposition into centromeres. Nature communications, 7, 13465.

49. Furuyama, T., Dalal, Y. and Henikoff, S. (2006) Chaperone-mediated assembly of centromeric chromatin in vitro. Proceedings of the National Academy of Sciences of the United States of America, 103, 6172–6177.

50. An, S., Kim, H. and Cho, U.S. (2015) Mis16 Independently Recognizes Histone H4 and the CENP-ACnp1-Specific Chaperone Scm3sp. J Mol Biol, 427, 3230–3240.

51. Yu, Z., Zhou, X., Wang, W., Deng, W., Fang, J., Hu, H., Wang, Z., Li, S., Cui, L., Shen, J. et al. (2015) Dynamic phosphorylation of CENP-A at Ser68 orchestrates its cell-cycle-dependent deposition at centromeres. Developmental cell, 32, 68–81.

52. Furuyama, T. and Henikoff, S. (2006) Biotin-tag affinity purification of a centromeric nucleosome assembly complex. Cell cycle, 5, 1269–1274.

53. Bobkov, G.O.M., Gilbert, N. and Heun, P. (2018) Centromere transcription allows CENP-A to transit from chromatin association to stable incorporation. The Journal of cell biology.

54. Subramanian, L., Toda, N.R., Rappsilber, J. and Allshire, R.C. (2014) Eic1 links Mis18 with the CCAN/Mis6/Ctf19 complex to promote CENP-A assembly. Open biology, 4, 140043.

55. Hayashi, T., Ebe, M., Nagao, K., Kokubu, A., Sajiki, K. and Yanagida, M. (2014) Schizosaccharomyces pombe centromere protein Mis19 links Mis16 and Mis18 to recruit CENP-A through interacting with NMD factors and the SWI/SNF complex. Genes Cells, 19, 541–554.

56. Nardi, I.K., Zasadzinska, E., Stellfox, M.E., Knippler, C.M. and Foltz, D.R. (2016) Licensing of Centromeric Chromatin Assembly through the Mis18alpha-Mis18beta Heterotetramer. Molecular cell, 61, 774–787.

57. Wang, J., Liu, X., Dou, Z., Chen, L., Jiang, H., Fu, C., Fu, G., Liu, D., Zhang, J., Zhu, T. et al. (2014) Mitotic regulator Mis18beta interacts with and specifies the centromeric assembly of molecular chaperone holliday junction recognition protein (HJURP). The Journal of biological chemistry, 289, 8326–8336.

58. Barnhart, M.C., Kuich, P.H., Stellfox, M.E., Ward, J.A., Bassett, E.A., Black, B.E. and Foltz, D.R. (2011) HJURP is a CENP-A chromatin assembly factor sufficient to form a functional de novo kinetochore. The Journal of cell biology, 194, 229–243.

59. Fachinetti, D., Logsdon, G.A., Abdullah, A., Selzer, E.B., Cleveland, D.W. and Black, B.E. (2017) CENP-A Modifications on Ser68 and Lys124 Are Dispensable for Establishment, Maintenance, and Long-Term Function of Human Centromeres. 40, 104–113.

60. Suter, B., Pogoutse, O., Guo, X., Krogan, N., Lewis, P., Greenblatt, J.F., Rine, J. and Emili, A. (2007) Association with the origin recognition complex suggests a novel role for histone acetyltransferase Hat1p/Hat2p. 5, 38.

61. Van Hooser, A.A., Ouspenski, II, Gregson, H.C., Starr, D.A., Yen, T.J., Goldberg, M.L., Yokomori, K., Earnshaw, W.C., Sullivan, K.F. and Brinkley, B.R. (2001) Specification of kinetochore-forming chromatin by the histone H3 variant CENP-A. Journal of cell science, 114, 3529–3542.

62. Bailey, A.O., Panchenko, T., Shabanowitz, J., Lehman, S.M., Bai, D.L., Hunt, D.F., Black, B.E. and Foltz, D.R. (2015) Identification of the posttranslational modifications present in centromeric chromatin. Molecular & cellular proteomics: MCP.

63. Hayashi, T., Fujita, Y., Iwasaki, O., Adachi, Y., Takahashi, K. and Yanagida, M. (2004) Mis16 and Mis18 are required for CENP-A loading and histone deacetylation at centromeres. Cell, 118, 715–729.

64. Li, Q., Zhou, H., Wurtele, H., Davies, B., Horazdovsky, B., Verreault, A. and Zhang, Z. (2008) Acetylation of histone H3 lysine 56 regulates replication-coupled nucleosome assembly. Cell, 134, 244–255.

65. Barski, A., Cuddapah, S., Cui, K., Roh, T.Y., Schones, D.E., Wang, Z., Wei, G., Chepelev, I. and Zhao, K. (2007) High-resolution profiling of histone methylations in the human genome. Cell, 129, 823–837.

66. Greer, E.L., Beese-Sims, S.E., Brookes, E., Spadafora, R., Zhu, Y., Rothbart, S.B., Aristizabal-Corrales, D., Chen, S., Badeaux, A.I., Jin, Q. et al. (2014) A Histone Methylation Network Regulates Transgenerational Epigenetic Memory in C. elegans. Cell reports.

67. Greer, E.L., Maures, T.J., Hauswirth, A.G., Green, E.M., Leeman, D.S., Maro, G.S., Han, S., Banko, M.R., Gozani, O. and Brunet, A. (2010) Members of the H3K4 trimethylation complex regulate lifespan in a germline-dependent manner in C. elegans. Nature, 466, 383–387.

68. Bessler, J.B., Andersen, E.C. and Villeneuve, A.M. (2010) Differential localization and independent acquisition of the H3K9me2 and H3K9me3 chromatin modifications in the Caenorhabditis elegans adult germ line. PLoS genetics, 6, e1000830.

