## Supplementary table&figure egends for "RbAp46/48^LIN-53^ and HAT-1 are required for initial CENP-A^HCP-3^ deposition and de novo centromere formation in *Caenorhabditis elegans* embryos"

Table S1. List of primers used for dsRNA production and genotyping.

| Targeted Gene | Forward/ Reverse | Sequence (5' to 3') | Reference |
|---|---|---|---|
| *hcp-3* | Forward | GCAAAATGAGAGCGTCACAA | (1) |
| | Reverse | TCAGAGATGTCGAAGGCAGA | |
| *knl-2* | Forward | TCGACTTGGTCGGACAGATT | (1) |
| | Reverse | TGCGATATGTGGCGTTATGT | |
| *lin-53* | Forward | CCCCGTTCTTGTACGATCTC | (1) |
| | Reverse | GGCAGATTGGTCTTCTCCAA | |
| *cbp-1* | Forward | ACAAAATCAACCATGGGGAGGTGCT | (2) |
| | Reverse | ACACTGTATCTTCAGTTGGTGGCGG | |
| *hat-1* | Forward | AGATGGAAGTCTG TGG CCA ACG TG | (2) |
| | Reverse | GGCGCAAAATTTCGCGTATGGGTAA | |
| *mys-1* | Forward | GCTTGCCAAACTTTTTCTGG | (2) |
| | Reverse | TTCTTTTTCGCAGCCTCAAT | |
| *mys-2* | Forward | CCCCAATTGAGCAGAAGCCTGGTAT | (2) |
| | Reverse | AAATCGATAAAAACAGAGGCGGCGG | |
| *lsy-12* | Forward | CCAAATCCTTCGGTCGAAA | This study |
| | Reverse | TTACCGTTATCGGAACCTCG | |
| *mys-4* | Forward | CCACGATATGCCAAAATTCC | This study |
| | Reverse | TTTTGAATTCCCAGAATCCG | |
| *set-1* | Forward | GATTTCCACCACCTGGAATGTTTAG | This study |
| | Reverse | GCATCTCTCTGCCCGCTTC | |
| *mcm-2* | Forward | TGGCTGATCGTGCTAACAAC | This study |
| | Reverse | ACGTTCAACTCGAATGGTCC | |
| *smc-4* | Forward | GGAAAACATTTGGTTGCCGTT | This study |
| | Reverse | AAACGGAAGAGTCGAGGGAT | |
| Seq-*hat-1* | Forward | TTTGCGATGAAAAATCTCACAACGAG | This study |
| | Reverse | CAAATTTACAGCAAATTTAGACGAATTTCGTG | |

Table S2. *C. elegans* strans used in this study.

| Strain name | Genotype | Reference |
|---|---|---|
| N2 | Wild-type | |
| OD421 | *unc-119 (ed3) III; ltSi4 [pOD833; hcp-3p/GFP::hcp-3; cb unc-119(+)] II; hcp-3(ok1892) III; ltIs37[pAA64; pie-1p/mCherry::his-58; unc-119 (+)]* | (3) |
| OD426 | ltIs37 [*pie-1p::mCherry::his-58*; *unc-119(+)*] IV. meIs1 [*pie-1p::GFP::LacI*]. | (4) |
| WYY34 | *meIs1 [pie-1p::GFP::LacI]; heSi45 [mcm-4::mCherry; unc-119(+)] IV* | This study |
| WYY44 | *hat-1(hkuSi10[gfp::hat-1]*) III | This study |

Figure S1. Deciphering how specific DNA structures used for microinjection affect the efficiency of new centromere formation by analyzing the frequency of AC segregation in 1-, 2-, 3-4-, 5-8-, 9-16-, 17-32-, 33-64-cell embryos. (A) HMW DNA arrays formed in the gonad after microinjection. The maximum projection of the entire gonad shows that HMW DNA arrays localize in the diplotene and diakinesis region. "*" indicates an artefate green dot outside the gonad tissue. Scale bar represents 5 µm. (B) Seperated channels of time-lapse images following a segregating AC in first mitosis in Figure 1. Scale bar represents 5 µm. (C) The schemetic of the p64xLacO for microinjection. To isolate circular, supercoiled DNA, p64xLacO was pre-treated with T5 exonuclease to degrade linear ssDNA, linear dsDNA and nicked plasmid DNA while preserving supercoiled DNA. For linear DNA, p64xLacO was digested by *AfaI*. (D) Representative images of nascent ACs in one-cell embryos, which were generated from microinjection of either supercoiled DNA (left) or linear DNA (right). White arrowheads indicate the ACs. * represents polar bodies. Scale bar represents 5 µm. (E) Quantification of the diameter of ACs (LacI::GFP dots) generated from injecting supercoiled DNA and linear DNA in one-cell embryos, based on measurement of the diameters of LacI::GFP dots in one-cell embryos that were indicating the size of ACs. The numbers of ACs (n) analyzed are shown. In the box plots, the boundary of the box closest to zero indicates the $25^{th}$ percentile, a line within the box marks the median, a black dot within the box marks the mean, and the boundary of the box farthest from zero indicates the $75^{th}$ percentile. Whiskers above and below the box indicate the 10th and 90th percentiles. Student's t-test was used to test the significance. ****: $p < 0.0001$. (F) Quantification of the segregation rate of ACs formed by injecting supercoiled DNA and linear DNA. AC segregation rates were scored as the % of cells with segregating ACs among all dividing cells containing ACs. ACs formed by supercoiled DNA have lower segregation competency than linear DNA at each cell stage. Although the segregation competency of all types of ACs improves over time, the difference is especially clear in early embryo stages. The number of cells (n) analyzed was indicated. A chi-square test was used to test significance. *$p < 0.05$, **$p < 0.01$ and ****$p < 0.0001$. NS means not significant. (Same data set as from Lin and Yuen, submitted back-to-back.) (G) The agarose gel image of sheared salmon sperm DNA (SS-DNA), with a mean size of about 6 kbs, was used for microinjection. (H) ACs with “complex” DNA context formed from microinjection of SS-DNA show positive HCP-3 signals in a 1-cell embryo, that expresses GFP::HCP-3 and mCherry::H2B. The yellow arrowhead indicates the position of the AC. Scale bar represents 5 µm.

Figure S2 (A) The inner centromeric protein, AIR-2, and the condensin II subunit, SMC-4, were both present on prometaphase ACs that were aligned on the metaphase plate. (B) Immunofluorescence signal of SMC-4 persisted on lagging ACs after anaphase. (C) SMC-4 persisted on the lagging endogenous chromosomes in a Hydroxyurea (HU)-treated embryo. A 12-µm line was drawn across the lagging chromosomes, and the signal intensities were measured. The plot profiles show signal intensities from each channel, Green: LacI; Red: SMC-4; Blue: DNA. Scale bar represents 5 µm. A higher-magnification view of the ACs (white square) is shown on the right, in which scale bar represents 2 µm. (D) Representative image of a one-cell embryos with a lagging AC in a strain that expresses GFP::LacI (green) and MCM-4::mCherry (red), from interphase to anaphase. The fold difference of signal intensity of MCM-4::mCherry on ACs and endogenous chromosomes is shown. The ACs that aligned at metaphase plates (white square) were used for calculating MCM-4::mCherry signal intensity. The yellow square represents the position of endogenous chromosomes, where MCM-4::mCherry is invisible. The signal intensity of MCM-4::mCherry on ACs and endogenous chromosomes were both subject to subtraction of the background signal (grey square). MCM-4::mCherry singanl intensity on ACs is about 20 fold higher than on endogenous chromosomes at metaphase, suggesting that DNA replication is still ongoing on ACs. (E) AC lagging at anaphase is not caused by LacI::GFP tethering. A nascent AC (white arrowhead) in an embryo expressing CENP-$A^{HCP-3}$::GFP showed the phenomenon of lagging chromatin during anaphase. (F) A representative image of a live cell (dashed line square) with an AC (white arrowhead), which segregated evenly in a multi-cell stage embryo. (G) The percentage of proper AC segregation among all ACs that were attempting to segregate in different embryonic stages in WT. (H) Representative live-cell image of ACs attempting to segregate in *mcm-2* RNAi-treated one-cell embryos. The time-lapse between the images was shown (mm:ss). Scale bar represents 5 μm. (I) Quantification of ACs segregation rates in WT and *mcm-2* RNAi-treated one-cell embryos. The number of samples (n) analyzed was indicated. Error bar indicates 95% C.I. of the SEM. No significant differences are found between WT and *mcm-2* RNAi-treated one-cell embryos, as analyzed by Fisher's exact test (NS, $p > 0.05$). (J) Immunofluorescence of BUB-1 on ACs in wild-type (WT) or *mcm-2* RNAi-treated one-cell stage embryos at prometaphase. Scale bar represents 5 μm. (K) The box plot shows the quantification of the normalized integrated density of CENP-A$^{HCP\text{-}3}$ signal on endogenous chromosomes and ACs in one-cell embryos.

Figure S3. Representative immunofluorescence images of H4K5ac, H4K12ac, H3K9ac, H4K20me, H3K4me, H3K4me2, H3K4me3, H3K56ac, H3K9me2, H3K9me3 and H3K27me3 on endogenous chromosomes and nascent ACs in one-cell embryos, with separated channels. Scale bar represents 5 μm.

Figure S4. (A) The expression of GFP::HAT-1 (WYY44) driven by the endogenous promoter in 4-cell embryos by live-cell imaging with no treatment and *hat-1* RNAi treatment. Scale bar represents 10 μm. (B) RT-qPCR is used to confirm the RNAi efficiency after microinjection of dsRNA of genes. Immunofluorescence of histone (C) H3 or (D) H4 on ACs in WT and *lin-53* RNAi-treated one-cell embryos. A higher-magnification view of the ACs (white square) is shown on the right. Scale bars in whole embryo diagrams and the magnified images represent 5 μm and 2 μm, respectively. A scatter plot shows the quantification result of the normalized integrated density of (C) H3 or (D) H4 signal on ACs in *lin-53* RNAi-treated one-cell embryos was compared with that in WT embryos. The number of ACs (n) analyzed was indicated. The integrated density of H3 or H4 was normalized to DAPI. Error bar indicates SD. Significant differences are analyzed by the student's t-test (**, $p < 0.01$).

Figure S5. Co-immunoprecipitation of GFP::HAT-1, RbAp46/48$^{LIN\text{-}53}$ and SMC-4 using embryo extracts from the transgenic strain expressing GFP::HAT-1. No antibody (Dynabead only) and Rabbit IgG immunoprecipitation (IP) was used as negative controls. Inputs and immunoprecipitated proteins were analyzed by Western blots.

Figure S6. (A) Immunofluorescence (IF) analysis of co-localization of CENP-A$^{HCP\text{-}3}$ and M18BP1$^{KNL\text{-}2}$ on nascent ACs in one-cell embryos. (B) The percentages of nascent ACs have both CENP-A$^{HCP\text{-}3}$ and M18BP1$^{KNL\text{-}2}$, have CENP-A$^{HCP\text{-}3}$ only, have M18BP1$^{KNL\text{-}2}$ only, and have neither CENP-A$^{HCP\text{-}3}$ nor M18BP1$^{KNL\text{-}2}$ were shown. n represents the number of cells analyzed.

### References


1. Lee, B.C., Lin, Z. and Yuen, K.W. (2016) RbAp46/48(LIN-53) Is Required for Holocentromere Assembly in Caenorhabditis elegans. *Cell reports*, **14**, 1819-1828.

2. Ho, V.W., Wong, M.K., An, X., Guan, D., Shao, J., Ng, H.C., Ren, X., He, K., Liao, J., Ang, Y. *et al.* (2015) Systems-level quantification of division timing reveals a common genetic architecture controlling asynchrony and fate asymmetry. *Mol Syst Biol*, **11**, 814.

3. Gassmann, R., Rechtsteiner, A., Yuen, K.W., Muroyama, A., Egelhofer, T., Gaydos, L., Barron, F., Maddox, P., Essex, A., Monen, J. *et al.* (2012) An inverse relationship to germline transcription defines centromeric chromatin in C. elegans. *Nature*, **484**, 534-537.

4. Yuen, K.W., Nabeshima, K., Oegema, K. and Desai, A. (2011) Rapid de novo centromere formation occurs independently of heterochromatin protein 1 in C. elegans embryos. *Current biology : CB*, **21**, 1800-1807.
