## Supplementary figures and images for "RbAp46/48^LIN-53^ and HAT-1 are required for initial CENP-A^HCP-3^ deposition and de novo centromere formation in *Caenorhabditis elegans* embryos"

### Supplementary figures

#### Figure S1

A

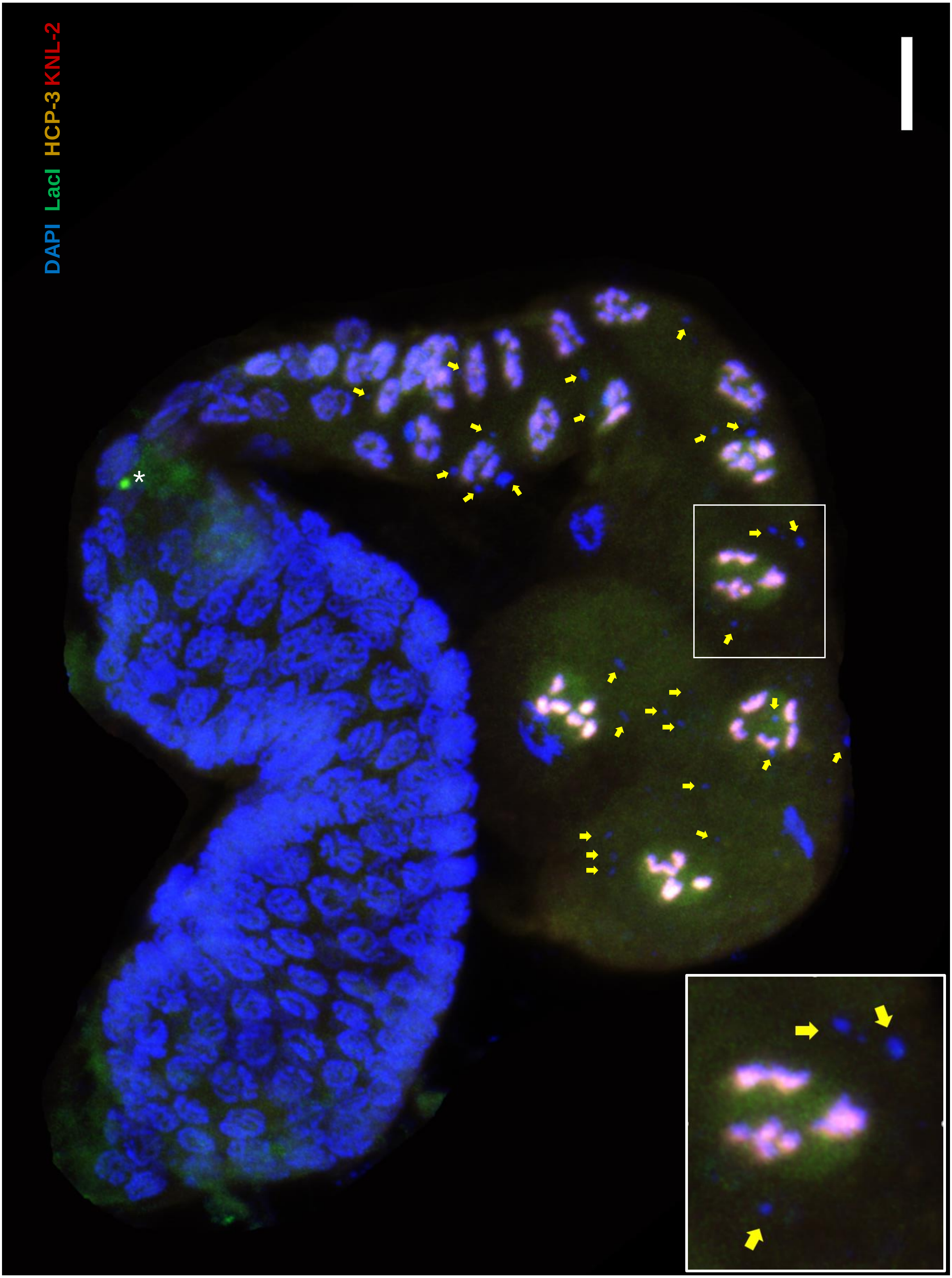

B

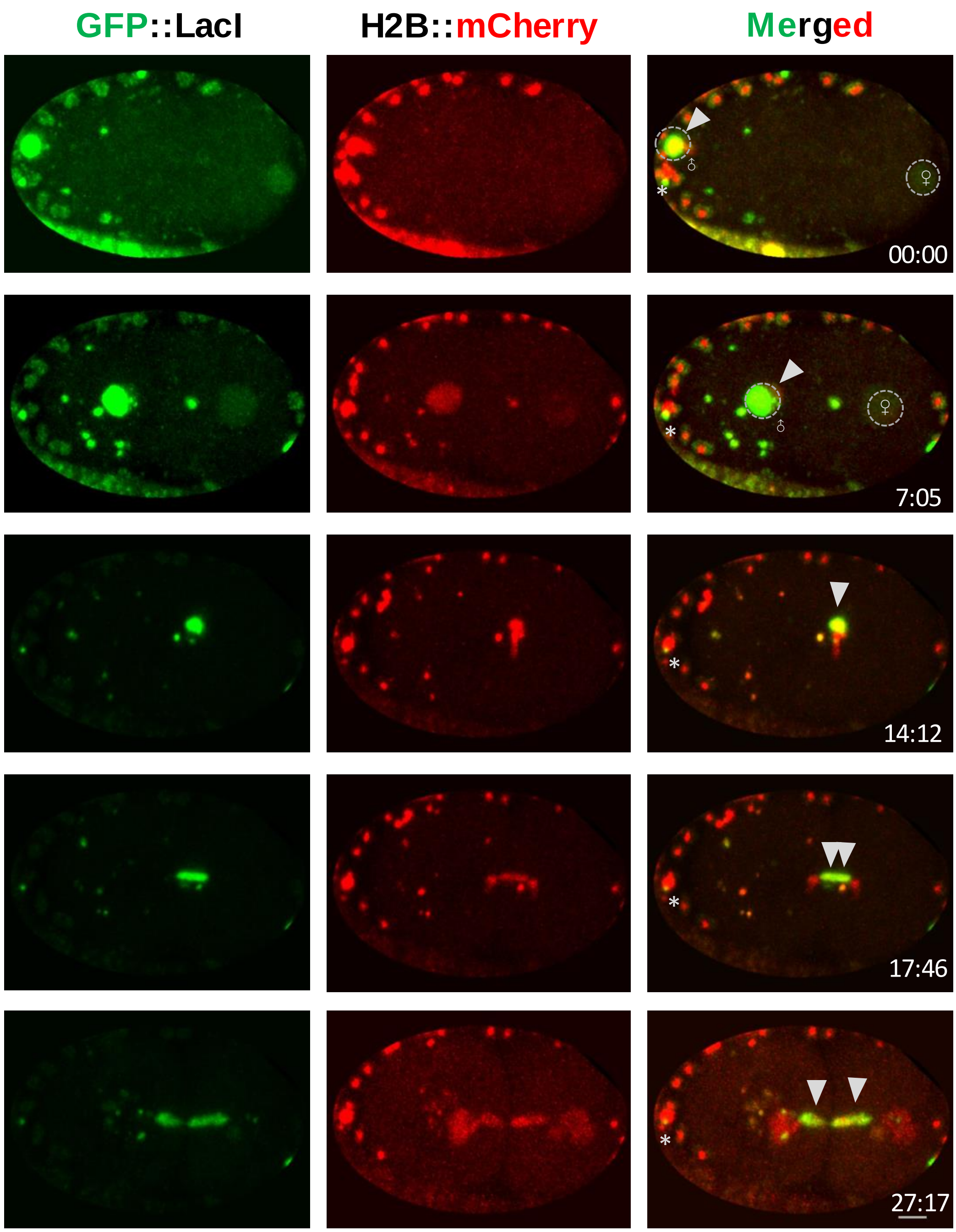

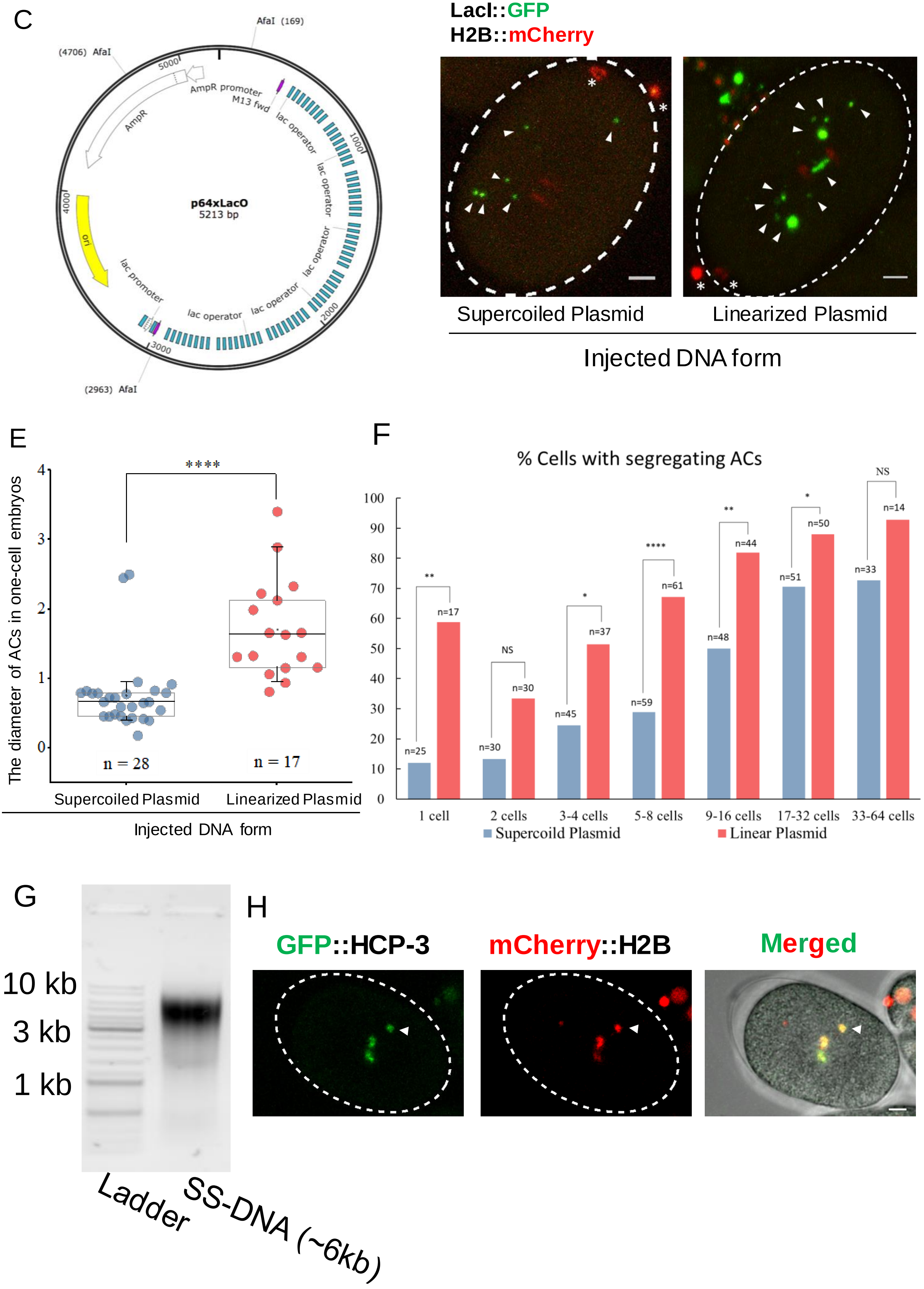

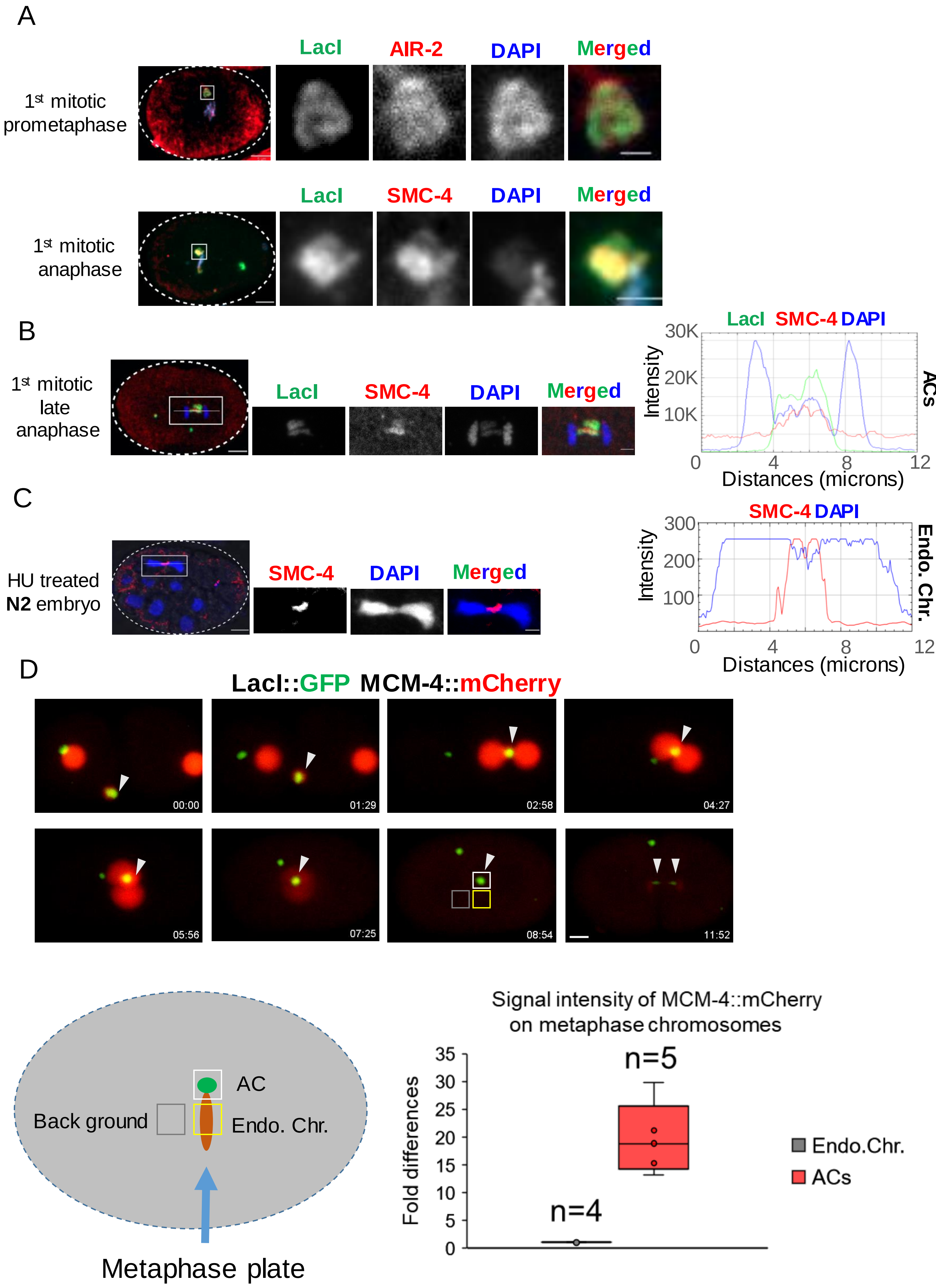

#### Figure S2

E

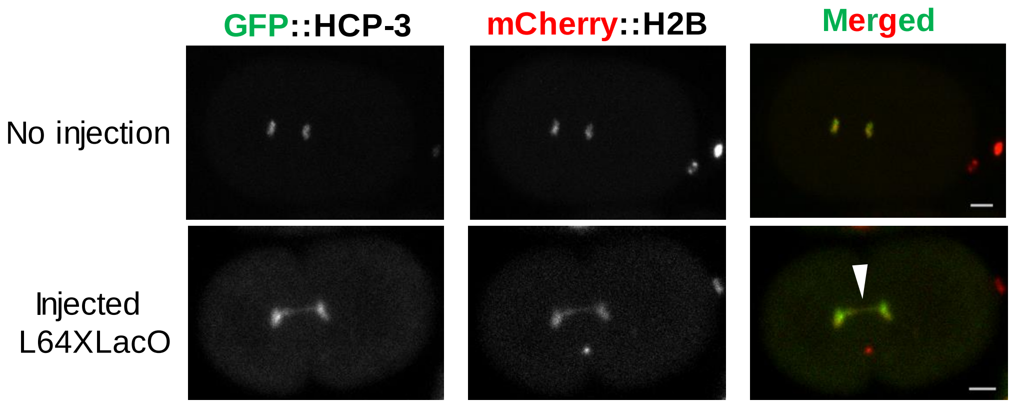


F

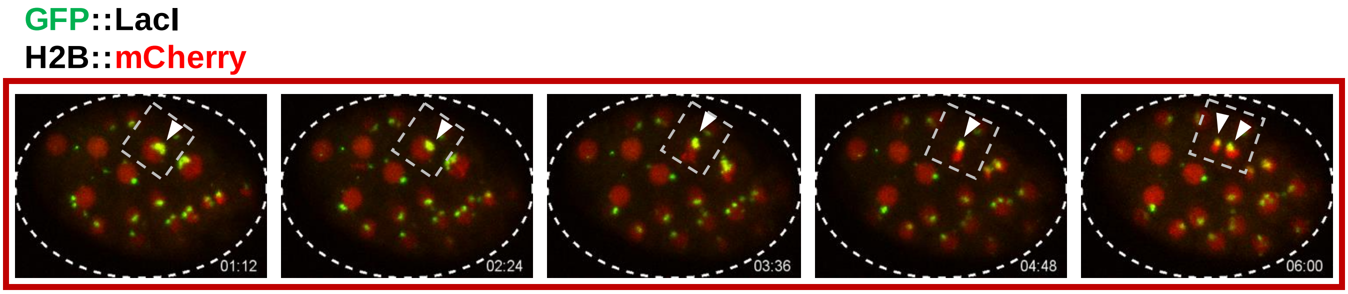


Proper AC segregation in multi-cell embryo

G

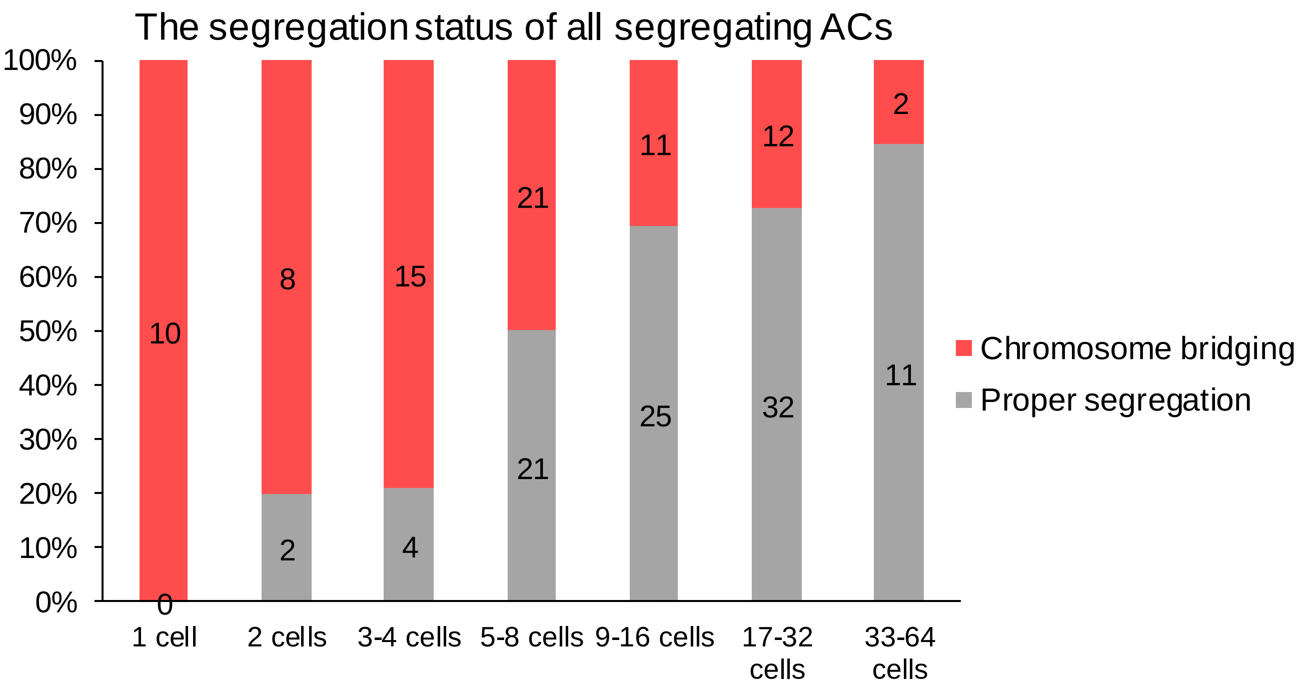

#### Figure S2

H

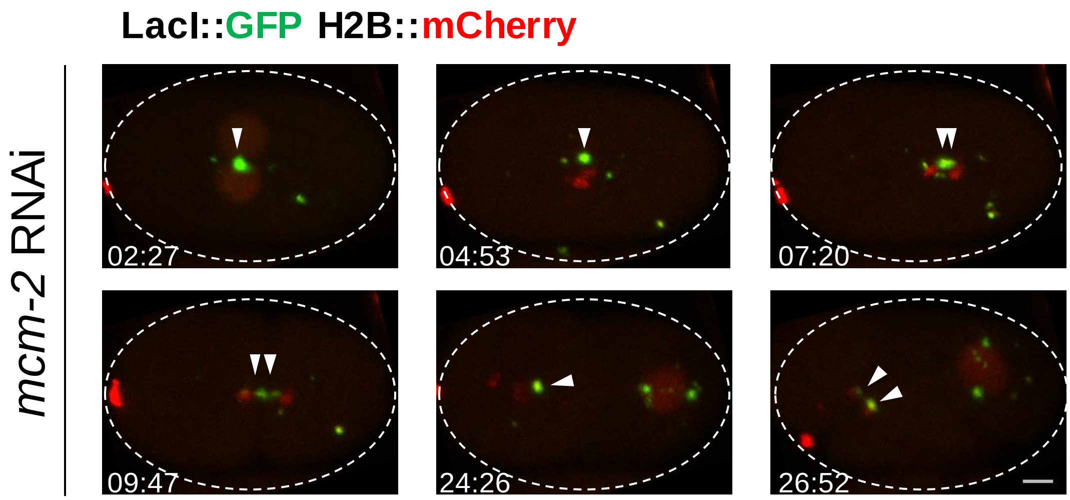


I

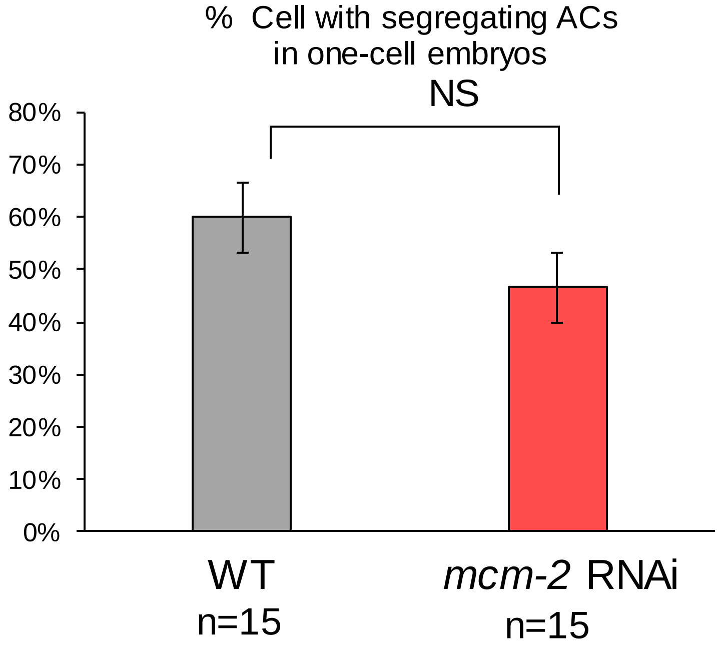


K

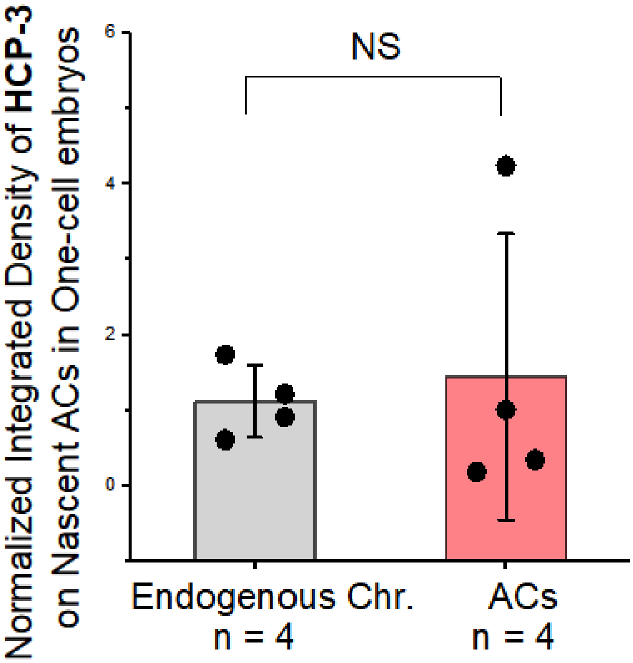


J

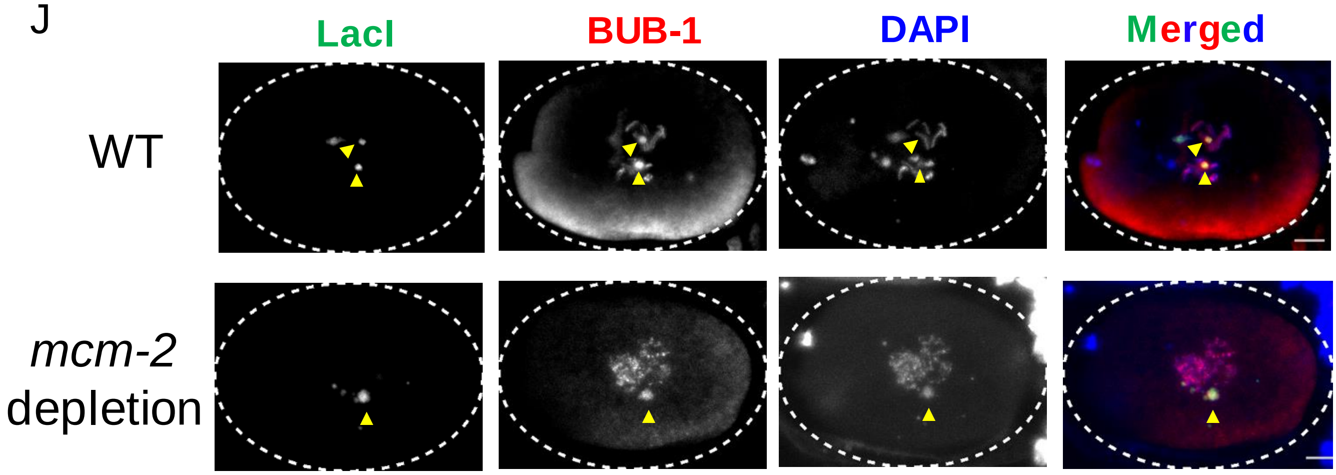

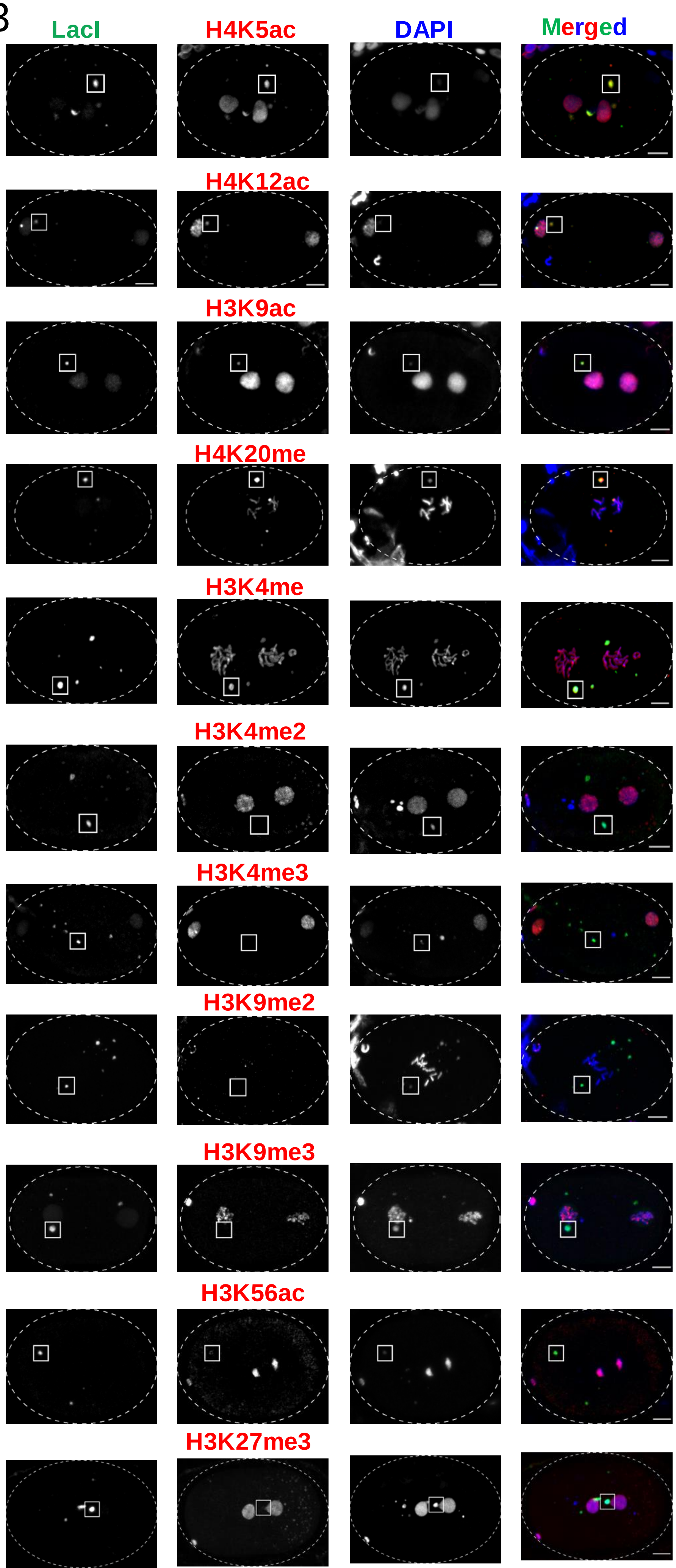

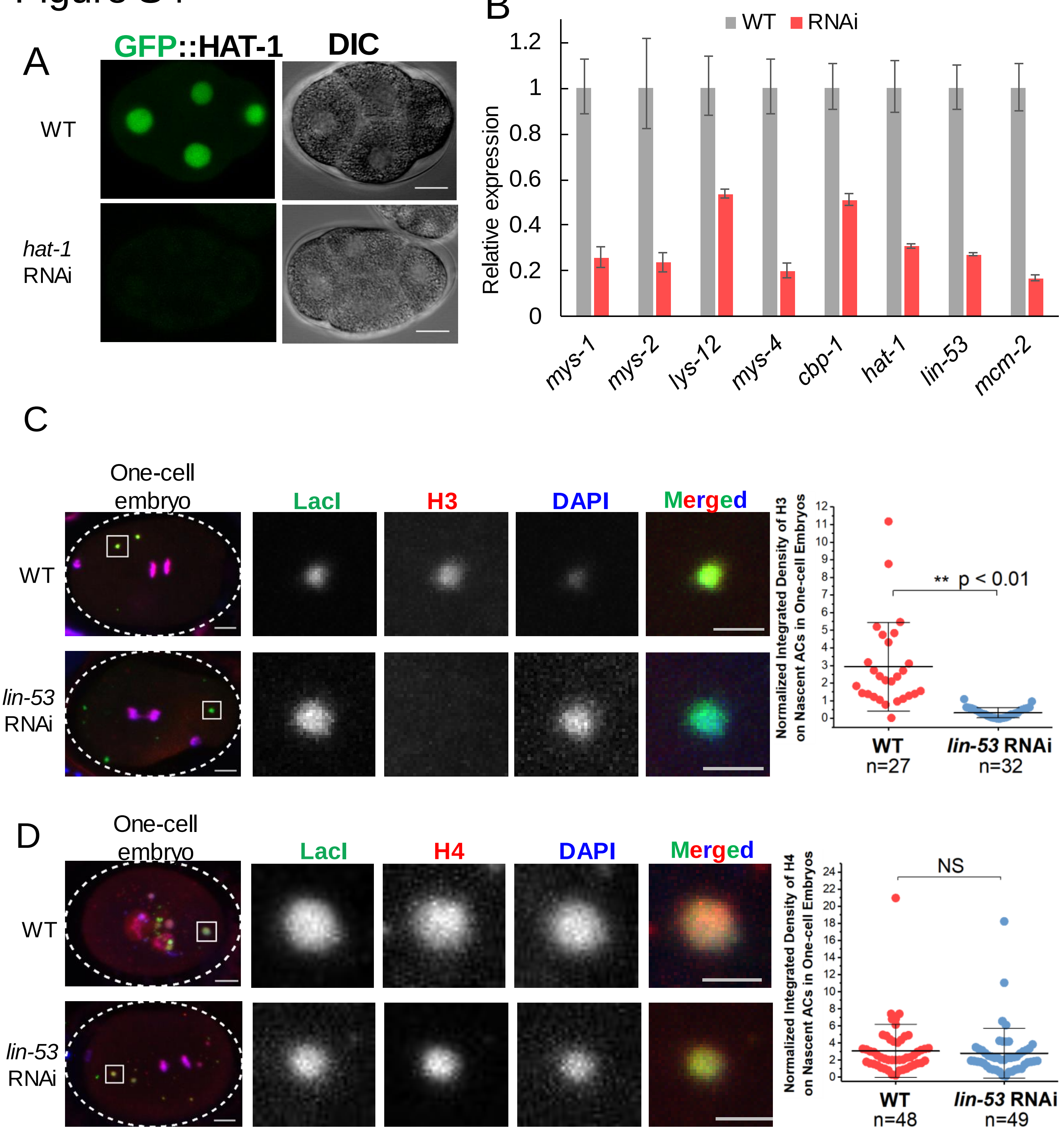

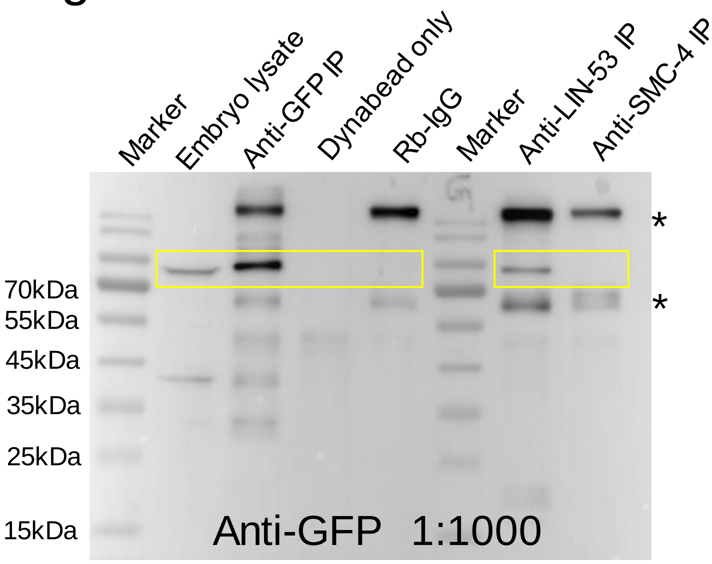

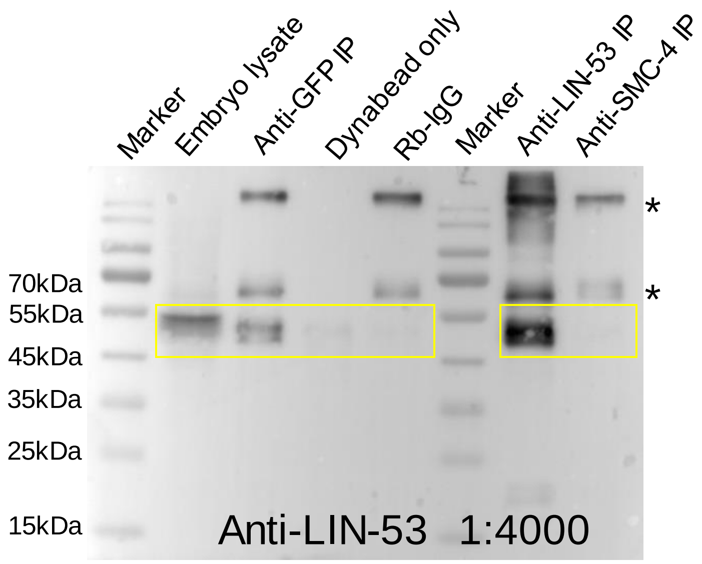

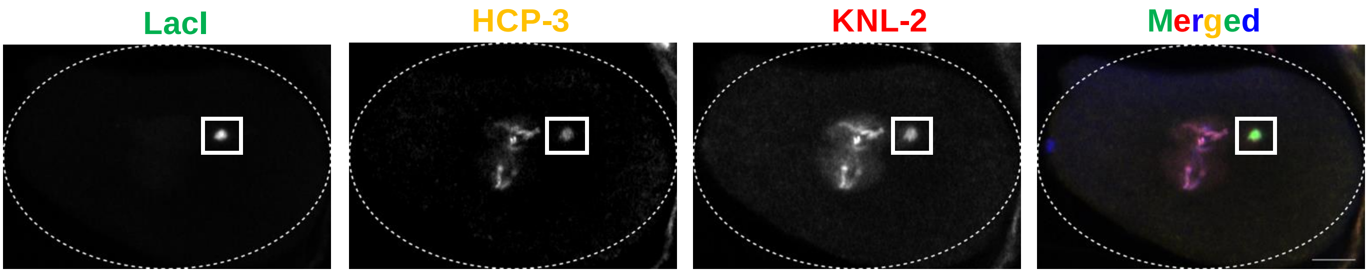

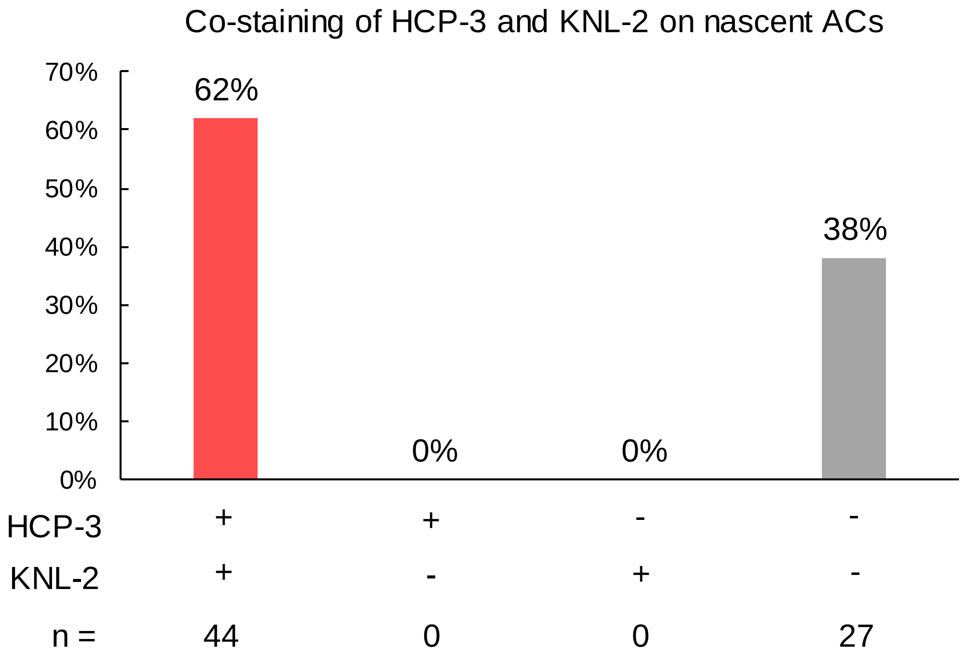
